# Temporal Transcriptome Analysis Uncovers Regulatory Modules Programming Embryo Development from Embryonic Morphogenesis to Post-Germination

**DOI:** 10.1101/2023.12.26.573376

**Authors:** Yen-Ching Wang, Wei-Hsun Hsieh, Chu-Jun Huang, Ya-Ting Jhan, Junpeng Zhan, Ching-Chun Chang, Tzung-Fu Hsieh, Jer-Young Lin

## Abstract

We profiled the soybean seed embryo transcriptome across embryonic development to post-germinative development to understand gene activities and regulatory networks promoting these processes. Transcriptomic landscapes feature highly prevalent transcripts which are categorized into early and late groups with major functions of reserve accumulation and energy generation, respectively, and both functions are dominant during late reserve accumulation as the transitioning stage. During the mid-reserve accumulation, regulatory events simultaneously dominate at the transcriptional and chromatin levels, followed by the emergence of distinct mRNA populations during late reserve accumulation throughout germination. We identified diverse functions conducted by sequentially activated genes across developmental stages. Gene coexpression network analysis reveals modules associated with developmental stages, some of which are enriched in genes with functions involved in specific developmental processes. We identified an early-desiccation-associated gene module, containing most transcription factors responsive to abiotic stress, within which one transcription factor is functionally validated to demonstrate increased drought tolerance in Arabidopsis overexpression lines. Finally, we found that a subset of genes is under purifying selection, surpasses the number of their Arabidopsis germination-specific homologs and most are active before germination from embryonic morphogenesis through dormancy, suggesting a potential role in governing physical dormancy in soybean compared to physiological dormancy in Arabidopsis. Our data represent a step toward identifying genes and regulatory networks in the soybean genome facilitating developmental programs across transition phases to bridge embryonic and germinative development.

## INTRODUCTION

Higher plant development can be divided into embryonic development, starting post-fertilization to initiate differentiation for axis and cotyledon formation, and post-germinative development, where differentiation recommences for adult growth. These two morphogenesis events are bridged by a period of developmental arrest, characterized by cessation of cell differentiation and division. During this period, several biological processes proceed subsequently for temporal transition, encompassing the accumulation of storage reserves, desiccation, dormancy, and germination (Fig 1). In the reserve accumulation and desiccation phase, cotyledon cells undergo enlargement and endoreduplication, facilitating the rapid accumulation of storage reserves, including seed-storage proteins and oil, to serve as energy sources during germination and seedling growth. Additionally, various compounds, such as disaccharides, are produced and may perform specific functions to protect against reactive oxygen species (ROS), stabilize protein conformation, modulate membrane integrity, and increase cell wall flexibility. Meanwhile, seeds acquire several important agricultural traits, including desiccation tolerance, seed longevity (the capacity to remain viable for long periods) (Leprince et al., 2017) and seed vigor (referring to germination rate, uniformity of seed germination, and tolerance to specific stresses post-germination) (Rajjou et al., 2012). Dormant seeds, the quiescent state, can survive in harsh environments and favorable conditions can break the dormancy for germination. Dormancy may result from exogenous factors (e.g., physical dormancy) or endogenous factors (e.g., physiological dormancy) (Baskin and Baskin, 2021). Soybeans exemplify physical dormancy with water-impermeable seed coat physically preventing water imbibition to delay germination and restrict radicle emergence, while *Arabidopsis* demonstrates physiological dormancy, where the absence of specific environmental cues (e.g., cold) inhibits germination. Germination progresses through three phases: Phase I involves rehydration, Phase II reactivates primary metabolism (e.g., nutrient reserve mobilization), and Phase III loosens the cell wall for radical protrusion, completing germination *sensu stricto* (Wang et al., 2015).

**Figure 1.**
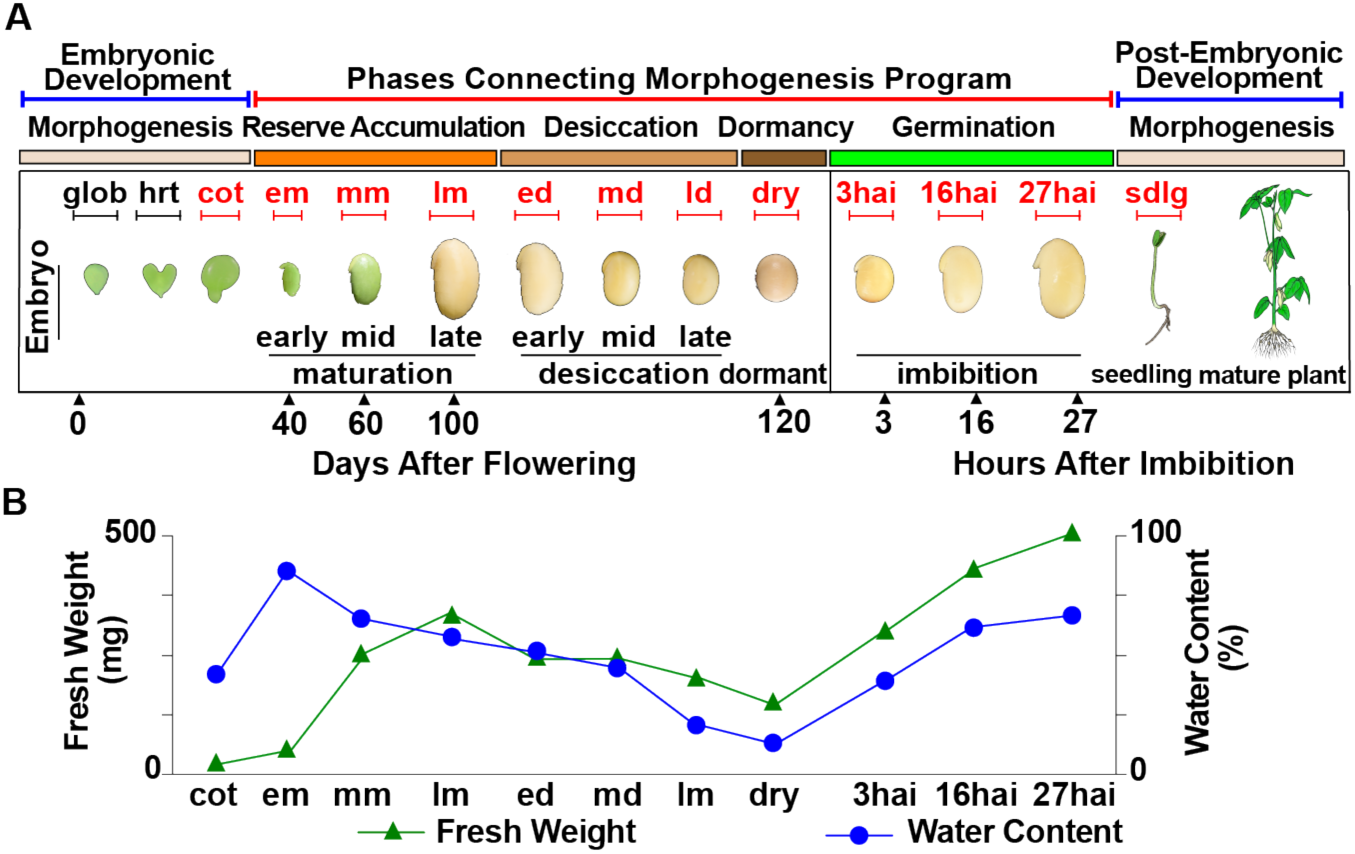
Schematic representation of soybean seed stages (A) and embryo weight (B).

To understand the molecular basis of seed development, transcriptome profiling has been applied to study temporal changes in genome-wide gene activities. These studies focus on various developmental aspects, such as early seed development to maturation (Le et al., 2010; Severin et al., 2010; Belmonte et al., 2013; Jones and Vodkin, 2013; Hofmann et al., 2019), late maturation (Verdier et al., 2013; Righetti et al., 2015; Pereira Lima et al., 2017) or germination (Buitink et al., 2006; Howell et al., 2009; Narsai et al., 2011; Dekkers et al., 2013). However, our knowledge of gene activities throughout the developmental stages connecting embryonic and post-germinative morphogenesis as a whole remains limited, especially in crops. Meanwhile, how genes and transcription factors (TFs) are organized into networks that govern biological events in each seed stage, ultimately guiding developmental transitions from embryonic to post-germinative morphogenesis, is relatively unknown.

Here, we conducted an in-depth time course transcriptome analysis between embryonic and post-germinative morphogenesis. Temporal mRNA populations across developmental stages are clustered into two groups, consisting of stages with green and yellow cotyledons, distinguished by the late reserve accumulation stage preceding desiccation. The transcriptomic profile throughout these phases is characterized by highly abundant mRNAs, comprising early and late expression groups that contain different major functions—reserve accumulation and energy generation, respectively, and shared functions, i.e., desiccation tolerance. The dominant functions conducted by up-regulated genes in each stage to promote developmental progress are revealed. During mid-reserve accumulation, regulatory events are significantly active at the transcriptional and epigenetic levels including genes associated with DNA methylation and histone modification, different from those enriched in germination, primarily in histone variants. In late reserve accumulation phase, various functions relative to desiccation tolerance are active, preparing for subsequent desiccation. We inferred gene regulatory networks associated with developmental stages and found an early-desiccation-associated module containing many TFs previously identified for coping with abiotic stress. The functional validation demonstrates that one TF predominantly expressed in this module may increase drought tolerance when this soybean TF is overexpressed in Arabidopsis. Finally, a substantial number of soybean genes experiencing purifying selection are active before germination, preceding the expression timeline of their Arabidopsis germination-specific homologs, which may contribute to the physical dormancy in soybean compared to the physiological dormancy in Arabidopsis. These results offer an atlas of gene activity that orchestrates developmental processes and reveals the regulatory modules associated with developmental stages promoting embryonic development to post-germinative development.

## RESULT

### Transcriptome Characteristics of Developing and Germinating soybean embryos

To obtain a comprehensive view of soybean transcriptome landscape between embryonic and post-germinative morphogenesis, we studied a detailed time-course of the transcriptome across six phases, including 12 developmental stages: (i) embryonic morphogenesis (cotyledon stage), (ii) storage reserve accumulation (early, mid-, and late maturation stage), (iii) desiccation (early, mid-, and late desiccation stage), (iv) dormancy, (v) germination (3hai, hour-after-imbibition, for germination phase I; 16hai for germination phase II; 27hai for germination III as radicle protrusion), and (vi) post-germinative morphogenesis (8-day seedling) (Fig. 1; Table 1). We profiled RNA-seq using soybean embryos from desiccation throughout germination (Supplemental File 1). We calculated normalized exonic reads from each replicate as fragments per kilobase of transcript per million mapped reads (FPKM) to perform pairwise correlation analysis, demonstrating high agreement among the three biological replicates, with correlation coefficients mostly ranging from 0.96 to 0.99 (Supplemental Fig. S1). We also obtained embryo transcriptomes from the Goldberg-Harada Gene Networks dataset (GSE99571), including embryonic morphogenesis, storage reserve accumulation, and post-germination, and observed a strong correlation among replicates after data processing (Supplemental Fig. S2). Together, RNA-Seq datasets used in our study exhibited high reproducibility, creating a comprehensive time course of embryonic developmental transcriptome for further analysis.

**Table 1.** Abbreviations of soybean development stage and postgermination.

| **Abbreviation** | **Description** |
| --- | --- |
| glob | Globular |
| cot | Cotyledon |
| em | Early maturation |
| mm | Mid maturation |
| lm | Late maturation |
| ed | Early desiccation |
| md | Mid desiccation |
| ld | Late desiccation |
| dry | Dry seed |
| 3hai | 3-hour after imbibition |
| 16hai | 16-hour after imbibition |
| 27hai | 27-hour after imbibition |
| sdlg | Whole seedling |

### Distinct mRNA populations emerge in late reserve accumulation phase throughout germination

To determine the temporal relationships of mRNA populations across different developmental stages, we conducted principal component analysis (PCA) and hierarchical clustering with correlation coefficients for all genes using FPKM (Fig. 2). Both analyses revealed strong similarities of the two group patterns: one group preceding desiccation and a distinctly separable group of stages succeeding desiccation. The clustering analysis dendrogram revealed two main clusters: one with green cotyledons and the other with yellow. The green cluster can be further divided into two sub-clusters: (i) before desiccation, including cot, em, and mm, and (ii) fully rehydrated, sdlg. The yellow cluster can be divided into two sub-clusters: (i) lm, representing the late reserve accumulation before desiccation, and (ii) stages experiencing desiccation (ed, md, and ld), dormancy, and rehydration (3hai, 16hai, and 27hai). The two-major-cluster pattern signified a substantial transcriptomic transition during embryo development, possibly attributed to extensive programmed water loss. Furthermore, lm was less associated with other stages that experience desiccation in the yellow cluster, suggesting a unique mRNA population in lm compared to stages from desiccation, dormancy, and germination. Together, the results indicate temporally controlled transition processes from embryonic to post-germinative morphogenesis, marked by changes in transcript population, initiating in the lm.

**Figure 2.**
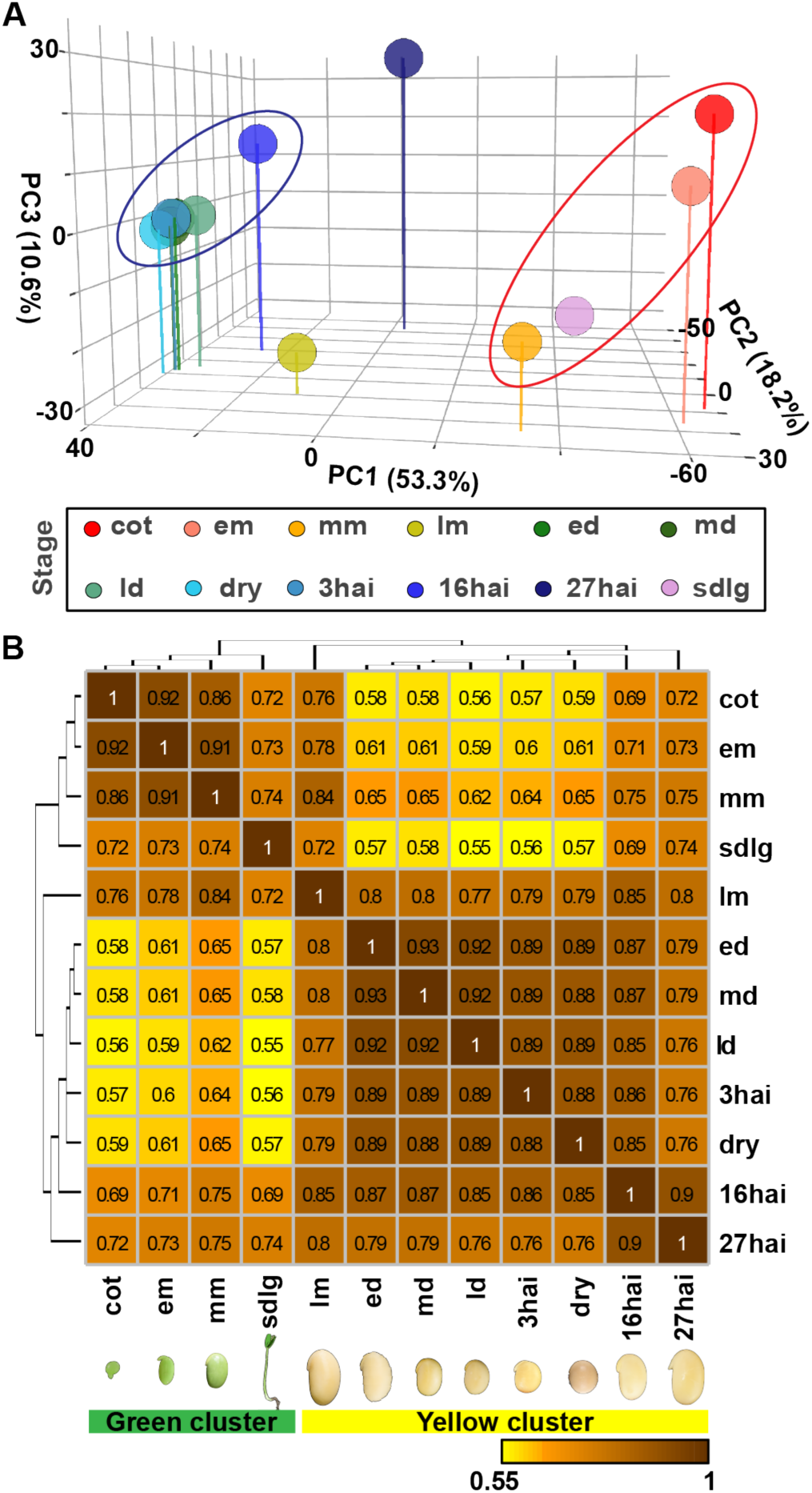
Relationship between the RNA populations across soybean embryo development. (A) Principal Component Analysis was conducted on transcripts from different embryo stages. Principal Components 1 to 3 (PC1 to PC3) together explained 82.1% of the variance in the mRNAs across the 12 stages. (B) Correlation coefficient analysis of RNA populations between stages was performed, and the resulting coefficients were hierarchically clustered to illustrate the association of developmental stages.

### Transcriptomic landscapes between embryonic and post-germinative morphogenesis are characterized by highly prevalent mRNAs

What are prevalent mRNAs in addition to transcripts encoding seed proteins (Goldberg et al., 1981) during seed development? We identified transcript abundance throughout development, primarily from em to 27hai, revealing a diverse range of prevalence (Fig. 3A). For instance, in mm, ∼40% of the total mRNA molecules was transcribed by eight genes, each of which produces mRNAs > 10,000 FPKM, defined as superabundant genes (Supplemental Table S2A). We identified 18 superabundant genes that were highly regulated with transcript levels fluctuating across development. Their expression patterns formed two groups: (i) the earlier superabundant gene group was highly active from em to mm consisting of seed proteins, including lectin (Le), glycinin (Gy1) and Kunitz trypsin inhibitor (KTi3); (ii) the subsequent group, highly active from lm to 27hai, comprised mitochondria and chloroplast proteins for energy production along with Late Embryogenesis Abundant (LEA) proteins primarily associated with desiccation tolerance.

**Figure 3.**
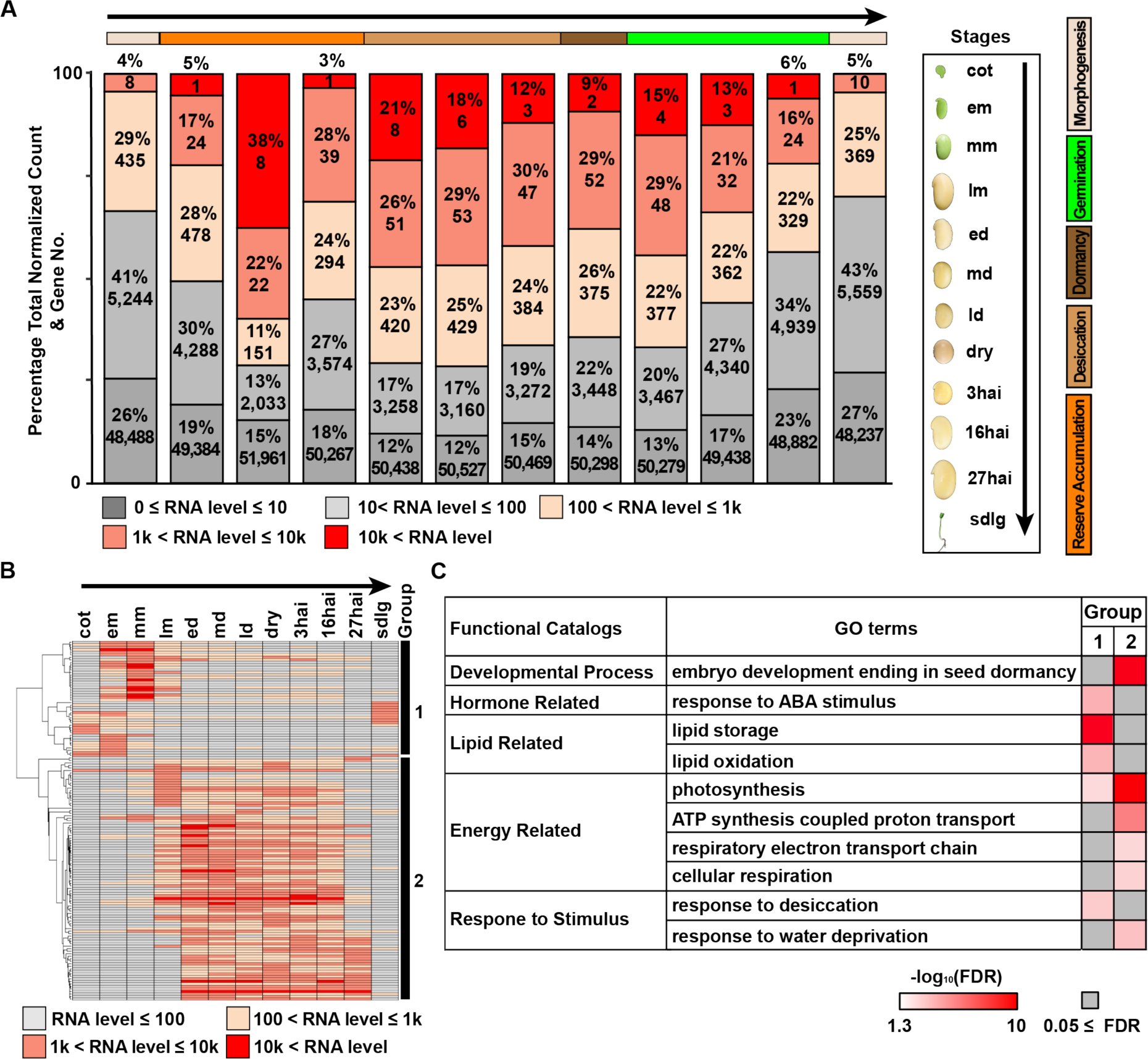
Highly prevalent transcripts across embryonic and post-germinative development. (A) The percentages of FPKM and the number of genes transcribing mRNA from varying prevalence levels are shown. (B) A heat map displaying the expression profiles of high-prevalence soybean genes (mRNA > 10,000 FPKM). The high-prevalent genes are hierarchically clustered into two groups, 1 and 2, based on mRNA levels across stages. (C) Biological process GO terms with FDR < 0.05 that are enriched in high-prevalence genes of clustering groups 1 and 2.

We then examined changes in the abundance fractions of high-prevalence genes, defined as genes with transcription levels > 1,000 FPKM to uncover 143 high-prevalence genes in which mRNA abundance increased from em, peaked in mm, and then dropped precipitously after 27hai (Fig. 3A). We conducted unsupervised clustering analysis to reveal two expression patterns similar to the superabundant genes: the first comprises 46 genes highly active from cot to lm, and the second includes 97 genes highly active from lm to 27hai (Fig. 3B). To characterize the major functions carried out in these two groups, we performed Gene Ontology (GO) analysis (FDR < 0.05) (Fig. 3C). We discovered common dominant functions shared by two groups of high-prevalence genes, including processes related to dormancy involved with ABA response, desiccation tolerance, and photosynthesis. Additionally, we identified specific functions in each group, such as lipid storage in the first group and ATP production in the second group. We further inspected the functions of lm-high-prevalence genes belonging to the two groups (Supplemental Table S2B). In cluster 1, the six high-prevalence genes were primarily seed proteins. The functions of the remaining 34 lm-high-prevalence genes in group 2 included desiccation tolerance (the major one, constituting 42% of lm-high-prevalence genes; Supplemental Fig. S3), salvage, raffinose biosynthesis, etc. Therefore, lm exhibits a distinctive high-prevalence mRNA profile, spanning two groups, each with different functions. Collectively, specific genes are highly transcribed from em to 27hai, with their expression patterns separated into early and late groups by lm, exhibiting major functions in storage reserve accumulation and energy generation, respectively, in addition to shared ones.

### Regulatory events at the transcriptional and chromatin levels concurrently dominate in the mid-reserve accumulation phase

When comparing consecutive developmental stages, significant quantitative changes can identify up-regulated genes at one stage compared to the previous stage, indicating the activation of genes to implement functions for programming development. Therefore, we focused on the activated genes by using DESeq2 (FDR < 0.05) and more than twofold increase to identify up-regulated genes in one stage compared to the earlier stage. Furthermore, we characterized the major functions carried out by up-regulated genes in one stage using Gene Ontology (GO) analysis (FDR < 0.01) (representative biological process GO terms in Fig. 4; the complete biological process GO terms in Supplemental Table S3; annotation of specific GO terms in Supplemental Table S4).

**Figure 4.**
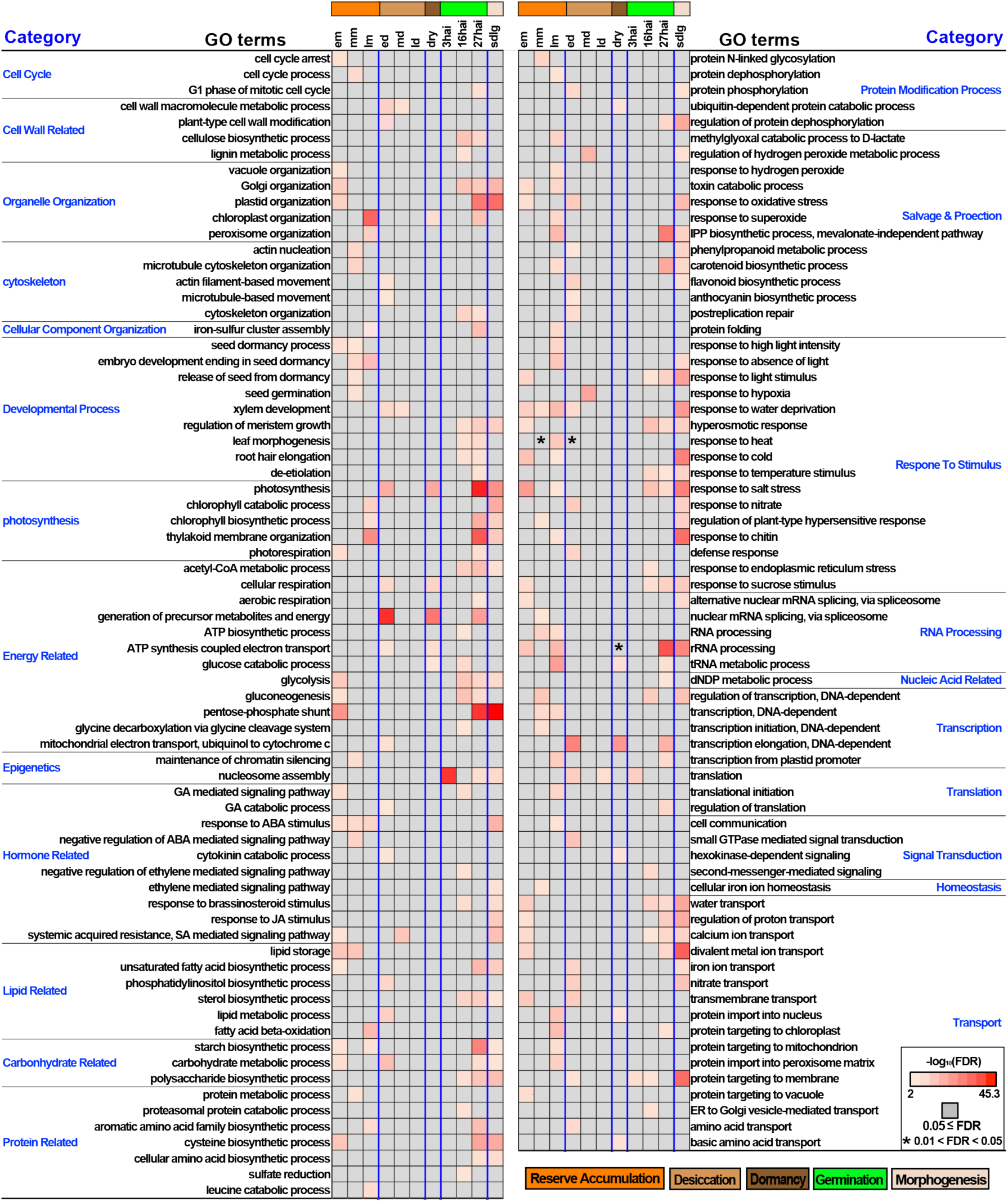
The biological process GO terms enriched in genes up-regulated than the previous stage. The representative GO terms with FDR < 0.01 are listed and the complete GO terms are listed in Supplementary Table S4.

We first asked whether transcriptional regulatory events are enriched in any stages. In mm, the “regulation of transcription” function was significant and 306 out of 446 genes in this GO term are TFs with diverse enriched functions (Supplemental Table S5). Meanwhile, epigenetics-relative functions, such as “maintenance of chromatin silencing” (Supplemental Table S6), were significant. In these GO terms, 17% and 41% of genes were associated with the two epigenetic hallmarks, DNA methylation and chromatin modification, respectively (Supplemental Fig. S4A). The DNA methylation-related genes included (i) DNA demethylase (e.g., *ROS1*), and, (ii) genes involved in *de novo* DNA methylation (RdDM pathway, e.g., *NRPD1B*, the largest subunit of PolV) (Supplemental Table S6A). Genes involved in chromatin modification were (i) chromatin writer, including H3K27 methyltransferase (e.g., *SWN*), H3K4 methyltransferase (e.g., *SUVH1*) and H3K9 methyltransferase (e.g., *SUVR5*), (ii) chromatin eraser, including histone deacetylase (e.g., SRT2), H3K27me3 demethylase (e.g., *REF6*) and H3K9 demethylase (e.g., IBM1), and, (iii) chromatin reader, including ATP-dependent chromatin remodeler (e.g., *BRM*) and transcriptional mediator (e.g., *MOM1*) (Supplemental Table S6B). The epigenetic-related function (i.e., nucleosome assembly) was significant again during germination but differed from those in mm, included genes up-regulated (i) in 3hai: histone variants, essential for programmed transitions (Jiang and Berger, 2023), e.g., *HIS1-3*, and (ii) 27hai: histone chaperones, e.g., *FAS2*. Together, regulatory events at the transcriptional and chromatin levels are active and coordinate to drive development in mm and distinct epigenetic processes underpin seed maturation and germination.

### Various functions are sequentially activated to facilitate the progression from embryonic morphogenesis to post-germinative morphogenesis

We examined significant functions, specifically the GO analysis results for up-regulated genes, to uncover dominant functions guiding development at each stage (Fig. 5 for representative GO terms indicated in this section; Supplemental Table S3 for complete GO terms; Supplemental Table S4 for annotation of up-regulated genes indicated in this section):

**Figure 5.**
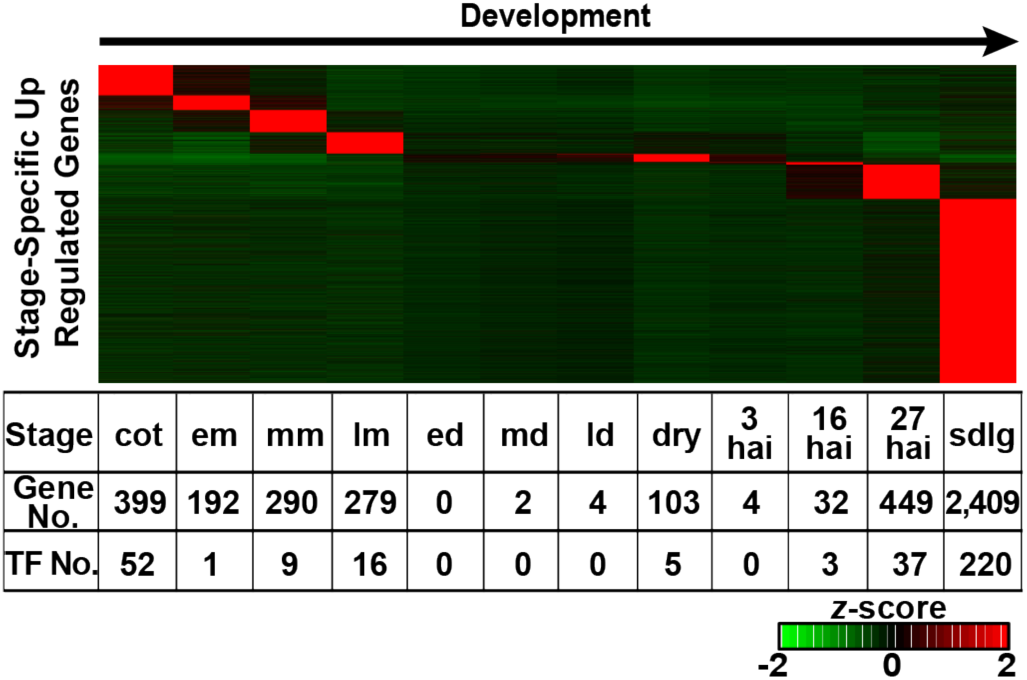
Expression profiles from soybean stage-specific genes during embryonic and post-germinative development. The scale, ranging from -2 (green) to +2 (red), indicates the relative number of standard deviations from the mean normalized RNA count for each stage-specific gene across all developmental stages.

#### Cell cycle

The functions of “cell cycle arrest” and “cell cycle process” were enriched in up-regulated genes in em and mm, including KRP7, negative cell division regulator (Cheng et al., 2013)) and CHR12, mediator of temporary growth arrest (Leeggangers et al., 2015), respectively. Up-regulated genes in 27hai were included in the function “G1 phase of the mitotic cell cycle”, which is critical for cell fate determination (Ma et al., 2015). The results suggest cell division ceases in em, the cell cycle pauses in mm, and then resumes for seedling growth in 27hai.

#### Cell wall

Functions related to cell wall remodeling were enriched beginning in ed, for instance, “plant-type cell wall modification” function including genes involved in (i) calcium-dependent cell wall loosening (e.g., *ANN5*) and (ii) remodeling biomechanical properties of the cell wall (i.e., Pectin Methylesterases, PMEs, like *CWINV2*). In 16hai, cell wall-related functions began to be dominant, such as “lignin metabolism process” (including gene *UGT72E1*) and “cellulose biosynthetic process” (including gene *CSLB04*). Together, cell wall remodeling functions are active upon desiccation underlying shrinking morphology and functions involved in cell wall synthesis are active in 16hai to support the following axis elongation and seedling growth.

#### Cytoskeleton

Relative functions were enriched in mm, ed and 27hai, possibly contributing to cotyledon enlargement during reserve accumulation, cotyledon shrinkage during desiccation, and preparation for radical protrusion in germination, respectively.

#### Cellular component organization

The “iron-sulfur cluster assembly” function was active in lm during chlorophyll degradation, suggesting the involvement of Fe-S proteins in processes related to dehydration, electron transfer shutdown, and disulfide reduction for ROS homeostasis in preparation for desiccation (Johnson et al., 2005).

#### Developmental process

Functions relative to dormancy were dominant throughout the reserve accumulation phase and included up-regulated genes involved in: (i) dormancy establishment/maintenance or inhibiting germination, such as *ASG5* (Bassel et al., 2011) and *HUB2* (Liu et al., 2007) in the “seed dormancy process” GO terms in em and mm, respectively; (ii) dormancy release or germination-induction, such as *ABH2* (Weng et al., 2016) in em-enriched function “seed dormancy process”. Additionally, in mm, “seed germination” function included nine genes inhibiting germination and 10 genes promoting it, e.g., *NAC040* (Supplemental Fig. S5A; Supplemental Table S7) (Song et al., 2022). Notably, *GmNAC040* was active from reserve accumulation throughout imbibition, whereas *AtNAC040* was active during germination only, highlighting their divergent expression patterns (Supplemental Fig. S5B). From 16hai to 27hai, dominant functions relative to development of diverse tissues (e.g., leaf and root) were enriched for autotrophic seedling formation. Together, the regulation of dormancy processes may be initiated during the early reserve accumulation phase, while a variety of development to form different tissues and cells are triggered during the mid-and late imbibition.

#### Photosynthesis

The function relative to chlorophyll degradation was enriched in lm when cotyledon turned yellow. Remarkably, in lm, ed and dry, functions relative to photosynthesis, such as “chloroplast organization”, “chlorophyll biosynthesis” and “thylakoid membrane organization” were enriched in up-regulated genes, including *HEMF1* (key enzyme in the biosynthesis of chlorophyll) and *VIPP1* (critical to maintaining thylakoid membrane integrity) (Gupta et al., 2021). In 16hai, the function “glycine decarboxylation via the glycine cleavage system” to process the toxic by-product generated in photosynthesis was enriched, preceding the 27hai-enriched function “photorespiration”. Together, although photosynthesis is shutting down in lm, specific mRNAs may be transcribed for storage in dry seeds, to be translated upon imbibition, facilitating rapid photosynthesis recovery in 27hai. Additionally, the salvage system for toxifying photosynthesis byproduct is established before photosynthesis resumes.

#### Energy

In ed, the “cellular respiration” function dominated, including *ADH1*, indicating fermentation under low oxygen conditions, and AOX1A, as alternative electron transport chain termination site to uncouple ATP synthesis. In dry, accumulated mRNAs were enriched with functions of energy precursor generation, e.g., “glucose catabolic processes”. In 16hai, the “ATP biosynthesis process” function and associated metabolic pathways became active, including “glycolysis”, “acetyl-CoA metabolic process”, and “gluconeogenesis” for acetyl-CoA production, connecting to the TCA cycle and the mobilization of stored reserves (e.g., lipid). In 27hai, “aerobic respiration” function was enriched, suggesting mitochondrial function restoration for O_2_ usage. Together, during desiccation, energy source includes fermentation and the electron transport chain shuts down. Transcripts involved in the upstream pathway of ATP generation are stored in dry seeds for translation upon imbibition, enabling rapid ATP production, and mitochondria activity resumes during the axis protrusion (27hai).

#### Hormone

Genes related to GA biosynthesis and signaling pathways were upregulated at em, while those involved in GA catabolism function were upregulated at ed, underlying decreased GA levels during late seed development. Diverse up-regulated genes enriched with ABA response functions were consecutively active during reserve accumulation. Cytokinin catabolism functions were enriched in ed, suggesting involvement in late embryo development (Tuan et al., 2019). In 16hai, functions related to GA, ethylene, and BR signaling pathways were enriched for germination and seedling growth.

#### Lipid

Lipid storage function became significant in em. Subsequently, beta-oxidation function was dominant in lm and ed, implying carbon partitioning from lipid breakdown to oligosaccharide in late seed development (Kambhampati et al., 2021). In ed, phosphatidylinositol biosynthesis was active, as signaling lipid for desiccation tolerance (Gasulla et al., 2013). From 16hai to seedling, functions related to the biosynthesis of sterols and fatty acids were dominant, serving as signaling hormones, precursors of brassinosteroids, and components of cell membranes for seedling growth. The results suggest soybean seeds store lipids early during reserve accumulation, and later during desiccation and germination, specific lipids contribute to progressing development.

#### Carbohydrate

Starch biosynthesis and plastid organization functions became active in em, which coincides with the conversion event of chloroplasts to amyloplasts (Borisjuk et al., 2005). In 16hai, “polysaccharide biosynthesis” function, included COB, the key regulator of directional cell elongation (Roudier et al., 2005), underlying radical protrusion. In 27hai, functions of “starch metabolic process”, “carbohydrate metabolic process”, and “maltose (degradative product of starch) metabolic process” were active, including genes involved in starch degradation, e.g., *PWD*. Together, soybean seeds may store starch early in reserve accumulation and its mobilization may occur during radical protrusion in 27ahi.

#### Protein and amino acids metabolism

In mm, the enrichment of the “protein metabolic process” function coincided with the transcription of superabundant seed protein genes. In lm, the catabolic process of the branched-chain amino acids leucine was dominant, potentially serving as alternative source for acetyl-CoA production (Gipson et al., 2017). In 16hai, functions related to protein mobilization, e.g., “proteasomal protein catabolic process”, were enriched. Meanwhile, assimilatory sulfate reduction, represented by the dominant function “sulfate reduction”, may subsequently lead to sulfite-containing amino acid biosynthesis, exemplified by the 27hai-significant “cysteine biosynthetic process” function which is involved in ROS response (Koprivova and Kopriva, 2014). Together, specific amino acids may serve as alternative energy sources during late reserve accumulation and desiccation phases. Protein mobilization may occur at 16hai, and, subsequently, specific amino acids may be involved in the ROS response.

#### Protein modification

In mm, the “protein glycosylation” function was enriched, potentially contributing to the glycosylation of the abundant seed protein, e.g., beta-conglycinin (Picariello et al., 2013). Furthermore, protein modification plays a role in signal transduction: (i) in lm, the enriched “protein dephosphorylation” function included genes encoding protein phosphatases, e.g., *ABI1*, involved in ABA signal transduction, and (ii) in ed, the “protein phosphorylation” function included kinase genes related to ABA signaling, e.g., *SnRK2.6*.

#### Salvage and protection

##### (i) scavenging toxic molecules

In lm, ROS response functions were enriched, such as “response to hydrogen peroxide”. Meanwhile, functions related to antioxidant production were significant, including the biosynthetic pathways of isopentenyl diphosphate (IPP) in lm and phenylpropanoid in ed, leading to the production of carotenoids, anthocyanins, and flavonoids. Additionally, in lm, the glyoxalase pathway may be involved in detoxification, as indicated by the significant “methylglyoxal (MG) catabolic process” function, and MG signaling may confer desiccation tolerance (Hoque et al., 2016). In 16hai, the reactivation of the IPP biosynthetic pathway may lead to the biosynthesis of chlorophyll and phytohormones (e.g., GA and auxin) for seedling growth.

##### (ii) repairing macromolecules

The “protein folding” and “postreplication repair” functions were dominant in lm and ed, respectively. Together, scavenging and repairing processes become active in lm prior to desiccation phase.

#### Response to stimulus

##### (i) light

In lm, most genes related to “response to absence of light”, e.g., *ETFQO*, an alternative substrate of the mitochondrial electron transport chain (Ishizaki et al., 2005), and those associated with “response to high light intensity”, e.g., light stress-response gene *SEP1* (Heddad and Adamska, 2000), were different. In 16hai before axis protrusion in 27hai, the “response to light stimulus” function included light-responsive genes such as *CRY2*, which is related to photomorphogenesis (Ponnu et al., 2019). Collectively, a complex light response occurs during chlorophyll degradation and mRNAs involved in light signaling are prepared for the upcoming light perception during axis protrusion.

##### (ii) hypoxia

The function “response to hypoxia” was enriched in md, suggesting a response to the absence of photosynthetic oxygen supply due to photosystem shutdown (Rolletschek et al., 2005).

##### (iii) abiotic stress

functions relative to abiotic stress responses were dominant in two periods: from em to dry and from 16hai to sdlg, suggesting a continuum in conferring desiccation tolerance and responses to rehydration.

##### (iv) nitrate

The functions “response to nitrate” and “nitrate transport” were enriched in ed and sdlg, including nitrate transporters. Because nitrate breaks dormancy (Hendricks and Taylorson, 1974), these results suggest that nitrate-relative genes, active in ed, may contribute to germination later, in addition to their role in nutrient transportation.

##### (v) Endoplasmic reticulum stress

Relative functions were active in 16hai, suggesting the accumulation of aberrant proteins and concurrent active protein repair activity during rehydration.

##### (vi) Sugar stimulus

Relative functions were active in em and from 16hai to sdlg, suggesting that sugar signaling triggers the accumulation of seed storage reserves and promotes seedling growth, respectively (Borisjuk et al., 1998; Baier et al., 2004).

#### RNA processing and nucleotide metabolism

During reserve accumulation, dominant mRNA processing may lead to extensive alternative splicing in the subsequent desiccation, potentially playing a role in the process (Srinivasan et al., 2016). In dry, enriched functions of rRNA and tRNA processes in highly accumulated mRNAs suggested a rapid recovery of the translation ability upon imbibition. Later, in 27hai, nucleotide metabolism-related functions were enriched, which may support seedling growth.

#### Transcription and translation

In mm, the function “transcription initiation” was significant, while in ed and dry, “transcription elongation” function was enriched. In 27hai, the major function “transcription from plastid promoter” co-occurs with chloroplast-related functions, e.g., thylakoid membrane organization. Transcripts relative to translation were enriched in lm and desiccation, along with dominant mRNAs involved in rRNA and tRNA processing in lm and dry. Together, transcription-related genes may contribute to the accumulation of high-prevalence mRNAs in mm; mRNAs involved in transcription and translation processes accumulate before germination, facilitating rapid recovery upon imbibition. Additionally, chloroplast activity resumes in 27hai during radical protrusion, consistent with the overall timing of recovering photorespiration and aerobic respiration.

#### Signal transduction and cell communication

In lm, the “cell communication” function was enriched, including genes involved in transport, hormone signaling pathways (e.g., ABA and ethylene), kinases and drought response. In dry, “hexokinase-dependent signaling” was enriched in accumulated mRNAs. In 16hai, the function “second-messenger-mediated signaling” was dominant. Collectively, lm-stage cells trigger internal changes through intercellular communication to transition from reserve accumulation to desiccation phase. Dry seeds store cascade regulatory signals, which later collaborate with extracellular signaling molecules from a suitable environment to promote germination.

#### Transport

##### (i) Calcium ion transport

Transport activity of calcium, acting as both a second messenger and a structural component that holds together cell walls (Thor, 2019), became active in 16hai, aligning with the significant functions in signal transduction and cell wall, as indicated above.

##### (ii) Transmembrane transport

Relative functions dominated in em and ed as a major pathway for secondary metabolite transport (Gani et al., 2021); the “protein targeting to membrane” function was significant in ed and 16hai, including ABA transporter ABCG40 and GA receptor GID1B. Collectively, the transmembrane transporters play a role in the developmental transitions.

##### (iii) Protein targeting to organelle

In em, “protein targeting to vacuole” was enriched, along with the significant “vacuole organization” function, indicating the development of protein storage vacuoles. In lm, functions related to protein targeting to organelles, e.g., chloroplasts and peroxisomes, were accompanied by significant functions of chloroplast and peroxisome organization. In dry, the major function “protein import into the nucleus” may coordinate with the dominant transcription-related function indicated above to resume nuclear function. The results suggest that protein targeting is involved in functional changes of the organelle, contributing to developmental transitions.

##### (iv) Amino acid/oligopeptide/protein transport

Transcripts enriched in “basic amino acid transport” were accumulated in dry seeds, aligning with their role as central metabolites in stress response and growth (Han et al., 2021).

##### (v) ER and Golgi transport

In 16hai, the dominant relative function coincided with the “response to ER stress”, suggesting active protein restoration and production during imbibition.

In summary, during the reserve accumulation phase, transcriptional activities are highly active in producing lipids and starch, followed by seed proteins; changes in organelle and cell wall structures occur accordingly and various functions are induced to prepare cells for the upcoming desiccation. Several functions unfold during late reserve accumulation and remain active throughout desiccation: chlorophyll degrades, the electron transfer chain shuts down via alternative electron acceptors, compounds and pathway alternative to Acetyl-CoA and aerobic respiration provide energy, specific metabolites serve as signal molecules involved in desiccation processes, and diverse functions contributing to desiccation tolerance, e.g., salvage and protection systems. Meanwhile, various mRNAs, including those involved in generating ATP, gradually accumulate and are stored in dry seeds to be translated upon imbibition. This process facilitates the recovery of cell activities and conveys signal cascades to initiate diverse regulatory networks for seedling growth, such as photomorphogenesis.

### Specific genes are preferentially active in distinct developmental stages

We determined genes that are active exclusively, or at elevated levels, at one developmental stage as stage-specific up-regulated genes (Fig. 5), defined as genes up-regulated (DESeq2 with an FDR < 0.05 and > two-fold increase) in one stage when compared to all other stages in pairwise comparisons. We found that the stages with the highest numbers of stage-specific up-regulated genes and TFs are sdlg and cot, representing the morphogenesis phase, with the fewest in the desiccation phase. Specific mRNAs were gradually transcribed to accumulate in dry seeds (Fig. 5; Supplemental Fig. S6A) and their enriched functions (Supplemental Fig. S6B) included: (i) protection from abiotic stress (e.g., response to hydrogen peroxide), (ii) accelerating the recovery of biological functions upon germination, including transcription-related functions (e.g., transcription-elongation) and energy-related functions (e.g., photosystem II assembly), and (iii) regulatory signal cascades (e.g., hexokinase-dependent signaling). Collectively, stage-specific up-regulated genes are preferentially active in specific stages, implying the involvement of organizing regulatory networks to program development. Additionally, dry seeds store specific mRNAs to protect cells and rapidly recover cell activities and regulatory signals upon imbibition.

### Identification of gene coexpression modules across seed embryo development

To characterize the sequential activation of gene regulatory processes throughout the development from embryonic and post-germinative morphogenesis, we identified coexpressed gene sets by employing weighted gene coexpression network analysis (WGCNA) on highly regulated genes. We included genes meeting stringent criteria (DESeq2 FDR < 0.01 and > four-fold increase) across developmental stages. A total of 36,383 genes were organized into 16 coexpression modules (M1 to M16), ranging from 64 to 6,926 genes and 11 to 662 TFs (Fig. 6, A and B). We analyzed the Z-scores of genes within each module across developmental stages to reveal coordinated gene expression patterns within each module (Supplemental Fig. S7).

**Fig. 6.**
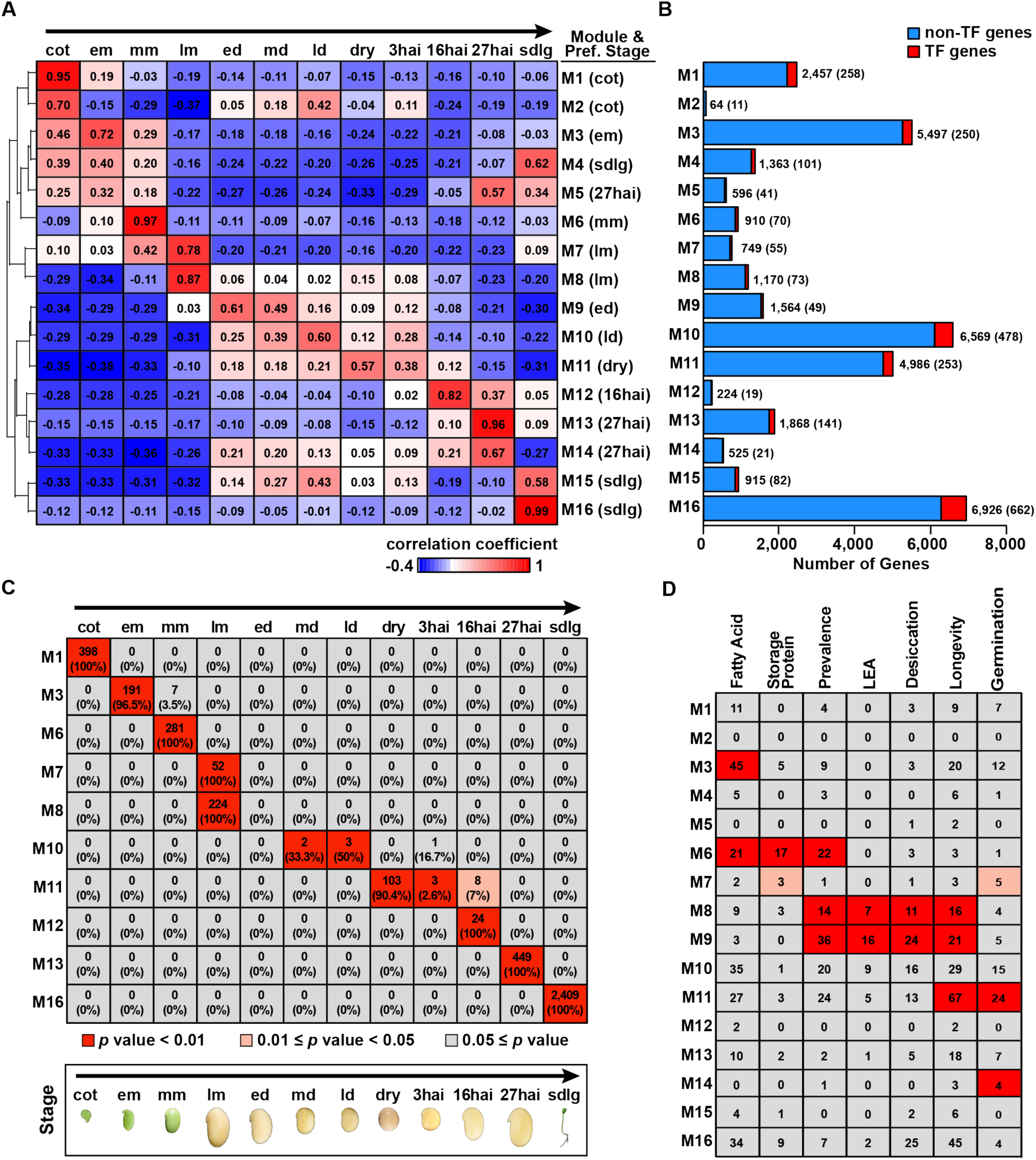
Gene modules identified by WGCNA. Heat map of the correlations between modules and developmental stages. The correlation coefficient analysis was performed using transcript levels in one module compared to developmental stages to obtain coefficients that were then hierarchically clustered to show the module-stage associations. Numbers of genes in each module. (C) Associations between the stage-specific gene sets and the modules. The heatmap illustrates *p*-values from hypergeometric tests for overrepresented genes in a pair of tests. The numbers in the boxes represent the overlapping gene counts and the proportions of these genes in the modules. (D) Associations of the gene sets with the coexpression modules. The heatmap shows *p*-values from hypergeometric tests assessing the overrepresentation of genes in a tested pair of gene sets.

Subsequently, we performed a permutation test to demonstrate that the average topological overlap of the 16 modules is significantly higher than that of randomly sampled modules (p-value < 0.005; Supplemental Table S8). Together, our analyses affirmed the reliability of the constructed regulatory modules.

### The modules are associated with temporal expression programs

To understand the association between coexpression modules and developmental stages, we calculated correlation coefficients, and then subjected the resulting r values to hierarchical clustering (Fig. 6A). The 16 coexpression modules were divided into two groups: module group 1, consisting of modules M1 to M7, which exhibited a relatively high correlation with stages from cot to lm; and module group 2, comprising modules M8 to M16, which was associated with stages from lm through desiccation to sdlg. Remarkably, lm was strongly associated with two modules: M7 signified the end of the reserve accumulation phase, while M8 marked the onset of desiccation. Therefore, lm encompasses two modules that bridge the distinct developmental phases of reserve accumulation and desiccation, aligning with the results of the high-prevalence gene analysis.

We asked whether modules are preferentially triggered in specific stages (Fig. 6A). Notably, the 16 coexpression modules can be categorized into two distinct types. First, six out of the 16 modules (M1, M6, M8, M12, M13, and M16) exhibited high correlations with specific stages (cot, mm, lm, 16hai, 27hai, and sdlg, respectively). Among them, the correlation coefficient for the module in this type was > 0.82 for one stage and < 0.37 for the other stages, suggesting a high degree of stage specificity for these modules. Second, the remaining 10 modules showed relatively strong associations with at least two stages, where one stage had higher correlation coefficients (0.78 to 0.57) indicating a preference for this stage and the second stage had lower correlation coefficients (0.49 to 0.21). For instance, M9 was primarily associated with ed and then md, while M10 was predominantly linked with ld and then md. These results indicate that the module performs functions contributing to the developmental program in one stage or is required across contiguous stages, representing a continuum. These two types of regulatory modules coordinate to advance embryonic morphogenesis to post-germinative development.

### The stage-associated modules are enriched with stage-specific up-regulated genes

We assessed the consistency between the assignment of genes to coexpression modules and the stage-based patterns of mRNA accumulation by examining the overlap between these modules and the previously identified stage-specific up-regulated gene sets (Fig. 6C). The results revealed that most modules contain up-regulated genes specific to one stage. For instance, M1, M3, M6, M12, M13, and M16 each contained up-regulated genes from a single stage. However, M10 included both md- and ld-specific up-regulated genes. Similarly, M11 included dry-, 3hai-, and 16hai-specific up-regulated genes. The overlap between each module and the corresponding stage-specific up-regulated gene set was greater than expected by chance (hypergeometric test p-value < 0.01) (Fig. 6C), indicating a specific subset of coexpressed genes within each module is associated with stage-specific functions. Together, these results suggest each stage is associated with one or more coexpression modules that trigger stage-specific gene regulatory processes, programming the development within one stage.

### Genes conducting specific functions are enriched in stage-associated modules

Regulatory modules with specific genes promoted diverse developmental processes. We employed a hypergeometric test to assess the relationship between temporal coexpression modules and genes linked to specific functions (Fig. 6D; Supplemental Table S9). We examined storage reserve genes to uncover that genes involved in fatty acid biosynthetic pathways were enriched in both em-associated-M3 and mm-associated-M6, and that genes encoding seed proteins were enriched in mm-associated-M6. Enriched genes in lm-associated-M7 and -M8 perform functions related to storage proteins and those associated with desiccation tolerance and longevity, respectively. Furthermore, lm-associated-M8 and ed-associated-M9 were enriched in (i) soybean LEA protein genes (Battaglia and Covarrubias, 2013), (ii) desiccation tolerance genes in legumes, and (iii) longevity genes in legumes (Detailed methods to identify genes with these functions were described in Supplemental File 1). We searched modules that significantly contained soybean genes homologous to Arabidopsis germination-specific genes (Supplemental File 1). These modules included 27hai-associated-M14 during germination, and, remarkably, modules preceding germination, such as lm-associated-M7 or modules with a relatively substantial number of genes, like em-associated-M3. Collectively, the results suggest that genes with specific functions are interconnected and organized into gene networks to facilitate specific biological processes to promote development.

### Early-desiccation-associated gene module harbors TFs involved in the abiotic stress response

Our module-stage association analysis indicated M9 is primarily linked to early desiccation and gene enrichment analysis revealed M9 is enriched in high-prevalence genes, LEA genes and genes relative to desiccation and longevity (Fig. 6, A and D). Therefore, M9 is most likely governed by TFs involved in conferring desiccation tolerance and longevity. The forty-nine TFs in M9 were associated with abiotic stress response (28 TFs, 57%), longevity (3 TFs), dormancy (1 TF, abscission and cell death (2 TFs), and 15 with other functions (Fig. 7, A and B; Supplemental Table S10). Remarkably, GmABI5 (the primary regulator contributing to desiccation tolerance, longevity, dormancy, and germination) (Zinsmeister et al., 2016) was in M9. In M9, GmABI5 was predicted to interconnect with genes involved in the biosynthetic pathway of Raffinose family of oligosaccharides (RFOs), from Myo-inositol to raffinose and stachyose (Supplemental Fig. S8). This prediction aligns with the fact that in Arabidopsis, ABI5 regulates the biosynthesis of RFOs, which is involved in seed maturity and vigor (Zinsmeister et al., 2016).

**Figure 7.**
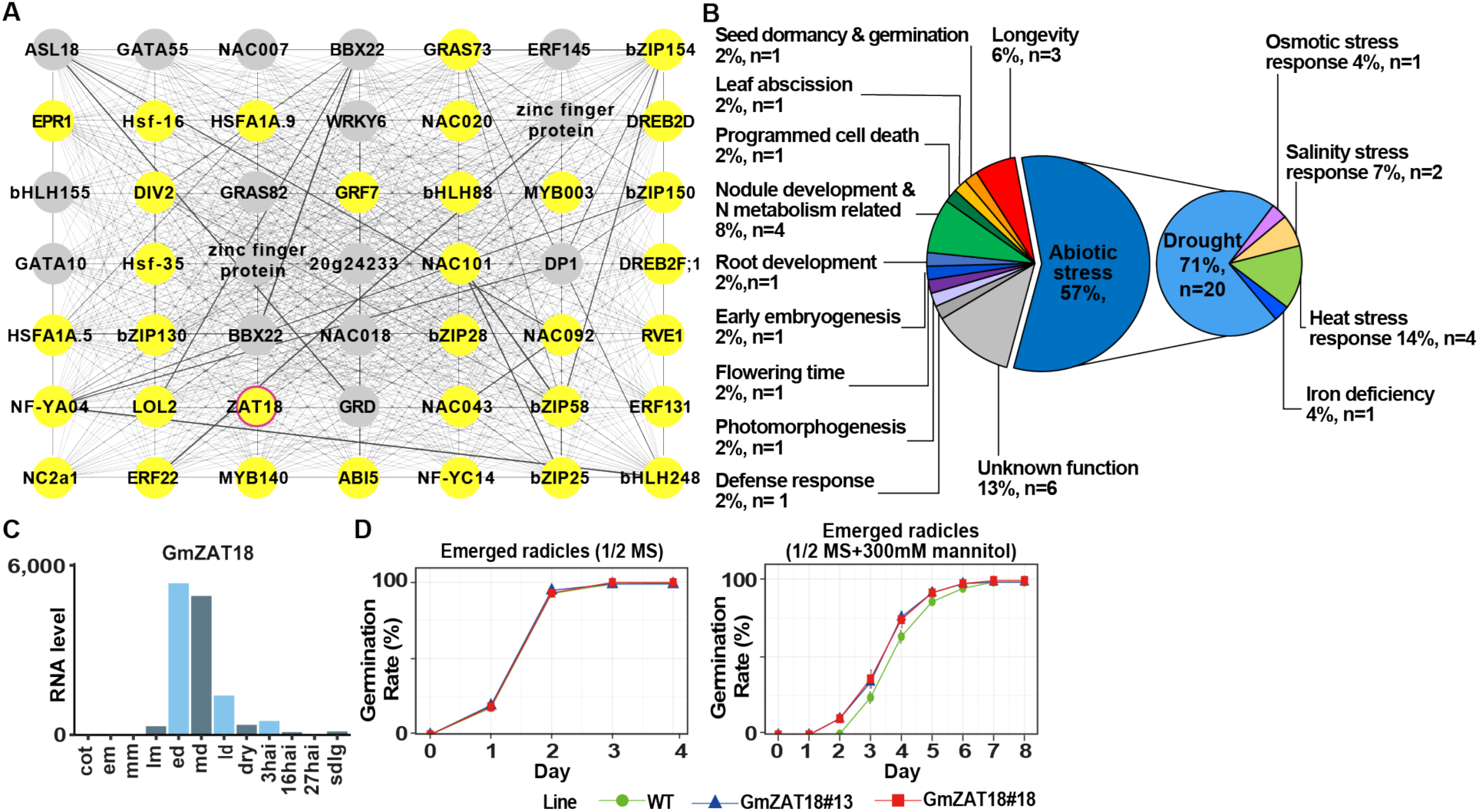
Regulatory module M9 and germination assay of GmZAT18 in Arabidopsis. A network of the 49 TFs visualized using Cytoscape (A), and their functional categories (B). (C) The expression pattern of GmZAT18 across developmental stages. The rates of emerged radicles are compared between overexpression lines and wild-type seedlings under control conditions (D) and in the presence of 300 mM mannitol (E) for four days.

Among the 49 TFs in M9, GmZAT18 (Glyma13g19560) was unique in that it became highly active in ed, maintained high activity in md, and then decreased precipitously in ld (Fig. 7C; Supplemental Fig. S9). Therefore, we deduced that GmZAT18 potentially contributes to desiccation tolerance. To test this prediction, we created GmZAT18 overexpression (OE) lines in Arabidopsis. GmZAT18 OE seeds had higher germination percentages than wild-type (WT) seeds under 300 mM mannitol treatment, with no difference under normal conditions (Fig. 7D; Supplemental Fig. S10). The data indicate drought tolerance in GmZAT18 OE lines is increased, which is similar to the enhanced drought tolerance observed in Arabidopsis with overexpressed AtZAT18, initially characterized as abiotic stress-induced genes in somatic tissues (Yin et al., 2017). Together, the data suggest the predicted module is reliable. Furthermore, we identified an early-desiccation-associated module that integrates TFs and genes conducting functions to program the desiccation process towards dormancy.

### Expression patterns of Arabidopsis germination-specific genes and soybean homologs are largely different

In the above analysis, we found several modules are associated with stages before germination and contain genes homologous to Arabidopsis genes highly active in germination (Fig. 6D). Similarly, the function “seed germination”, dominant in reserve accumulation phase, included upregulated *GmNAC040*, which is homologous to the Arabidopsis germination-specific genes *AtNAC040* (Supplemental Fig. S5). This raises the question of how many genes in the soybean genome exhibit different temporal expression patterns compared to their Arabidopsis homologs that are involved in germination. Previous microarray studies defined germination-specific genes in Arabidopsis (Narsai et al., 2011), of which we focused on 137 genes with mutant phenotypes and/or known sub-cellular localization (details in Supplemental file 1). The RNA-Seq profiles of these 137 Arabidopsis genes were retrieved from a public database for hierarchical clustering analysis, revealing that 49 of the 137 Arabidopsis genes were highly active during germination (Fig. 8; Supplemental Fig. S11). These 49 Arabidopsis have a higher number of soybean homologs, 129 soybean genes. Except for one soybean gene remaining inactive throughout seed development and post-germination, the mRNAs of the remaining 126 genes were hierarchically clustered into three groups showing distinct expression patterns: only 10 genes (group B in Fig. 8C) were highly active during germination, similar to their Arabidopsis homologs, while the remaining 116 genes were highly active either from lm to 16hai (group A) or from cot to lm (group C). Together, a subset of soybean genes exhibits a larger copy number and distinct expression patterns compared to their homologous Arabidopsis germination-specific genes.

**Figure 8.**
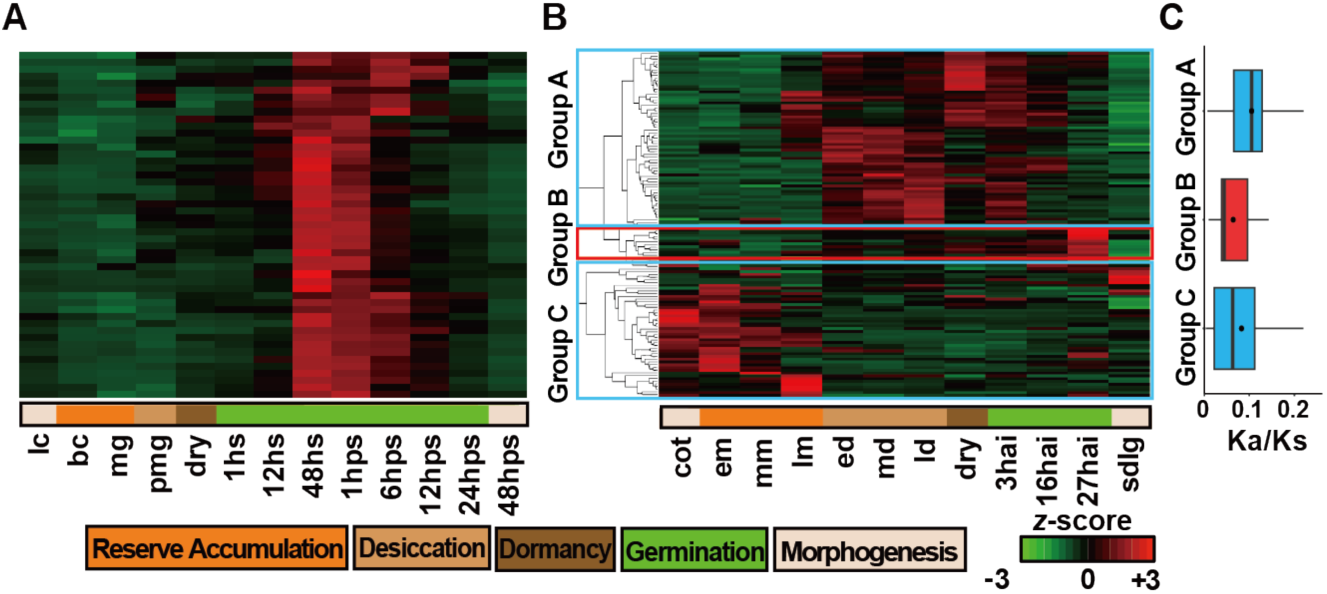
Gene expression pattern of the Arabidopsis germination-specific genes and their soybean homologs. (A) Heatmap of gene expression for the 49 Arabidopsis germination-specific genes during seed development through post-germination. (B) Gene expression heatmap of soybean homologs of the Arabidopsis germination-specific genes were hierarchically clustered into groups A and C, representing genes active before germination. Group B denoting genes active during germination. (C) Selection pressure analysis indicated by Ka/Ks values. The scale, ranging from - 3 (green) to +3 (red), indicates the relative number of standard deviations from the mean normalized RNA count for each gene across all developmental stages. Linear cot, lc; bent cot, bc; mg, post mature green; dry, dry seed; hour of stratification, hps; hour post-stratification.

### Divergent copy numbers and expression patterns among the homologous counterparts of soybean genes and Arabidopsis germination-specific genes

To further reveal expression divergence among soybean homologs corresponding to each *Arabidopsis* gene, we examined the expression patterns of soybean homologs. The 49 Arabidopsis genes were divided into two groups based on their soybean homolog numbers (Supplemental Table 11, A and B). In group 1 (six Arabidopsis genes and six soybean homologs), each Arabidopsis gene has one soybean homolog; none of them exhibits similar expression patterns between Arabidopsis genes and their soybean homologs. In group 2 (43 Arabidopsis genes and 120 soybean homologs), each Arabidopsis gene has multiple soybean homologs from two to seven. Soybean homologs exhibit either the same expression pattern (in the same expression group) or the combination of two of the expression patterns (in two expression groups). For instance, all four soybean homologs of AT1G68990 were in expression group A, while AT5G50760 has five soybean homologs, with two in group B and three in group C (Supplemental Table 11A). Together, the results suggest that duplication events occurred frequently and expression patterns evolved rapidly after soybean diverged from Arabidopsis ∼90 million years ago (MYA) (Grant et al., 2000).

### Widespread purifying selection at soybean genes homologous to Arabidopsis germination-specific genes

Given that the soybean genome has undergone at least two rounds of whole genome duplication (WGD) (Lin et al., 2010), we investigated the origins of the multiple soybean homologs (Supplemental Table S11C). For the three expression groups of soybean homologs, 74-82% originated from the recent WGD-I (∼13 MYA), while 11-22% were from a more ancient WGD-II (∼59 MYA), which percentages were not significantly different from all genes in the soybean genome (chi-square test, p < 0.05). These results suggest that the duplication events in these genes are closely linked to the history of soybean WGD events, rather than being more frequent in either ancient or recent duplications. To understand the selective pressures shaping the fate of gene activities, we examined the sequence divergence. The Ka/Ks ratios were less than 1 in all three groups (0.105±0.063, 0.064±0.044 and 0.083±0.072 for group A, B and C, respectively), indicating that purifying selection continuously eliminates deleterious mutations at these loci. Therefore, these soybean genes may contribute to the survival of the population, despite exhibiting different expression patterns from their homologous counterparts in Arabidopsis germination-specific genes.

Together, these results may suggest one of the molecular bases for the distinct dormancy types, with physical dormancy in soybean compared to physiological dormancy in Arabidopsis, for which we have developed a model (Fig. 9). A considerable number of soybean genes, homologous to relatively few Arabidopsis germination-specific genes, are active before seed germination, which has the additive genetic effect of preventing the establishment of physiological inhibitory mechanisms and, therefore, contributing to physical dormancy in soybean embryos. While we cannot completely rule out the possibility that they have undergone evolutionarily neofunctionalization resulting in displaying different functions before germination, we prefer this model because some of the substantial soybean genes may retain the same biological function as their Arabidopsis counterparts associated with germination-related duty to avoid physiological dormancy in soybean.

**Figure 9.**
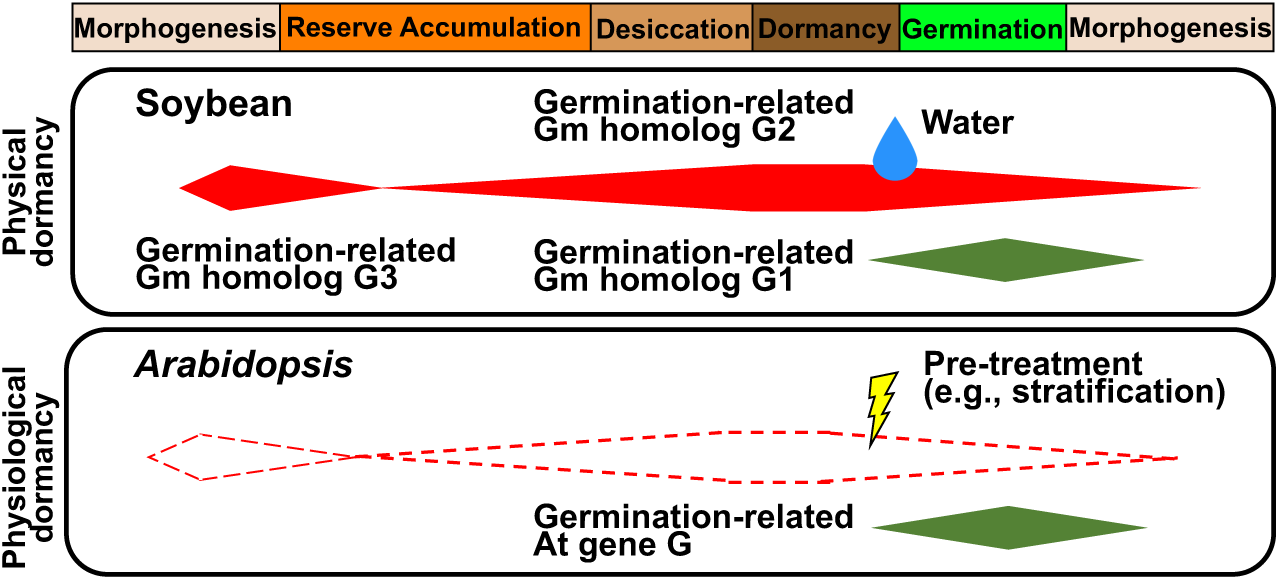
The model depicts physical dormancy in soybean and physiological dormancy in Arabidopsis. The green and red solid rhombi represent gene activities, while the red dot rhombi indicate the absence of Arabidopsis homologous counterparts for duplicated genes in soybean.

## DISCUSSION

Embryos prepare for desiccation in em by activating genes involved in desiccation tolerance. Meanwhile, lipid and starch biosynthesis genes become active, followed by seed protein genes active in mm. Remarkably, in mm, both regulatory mechanisms at transcriptional and epigenetic levels are dominant, which may contribute to reserve accumulation in the current stage and the biological process in the following stages. It is known that epigenetic mechanisms like histone modifications are involved in activating seed protein and fatty acid biosynthesis genes in *Phaseolus* and *Arabidopsis* seeds (Sundaram et al., 2013; Huang et al., 2022). What genes in the soybean genome are regulated directly at the epigenetic level in mm still remains largely unknown.

Transcriptome profiles in lm are unique as compared to stages before and after lm in several ways. First, transcriptome association analysis results show that it acts as an outgroup compared to other stages in the yellow-cotyledon cluster (Fig. 2). Second, overrepresented mRNAs exhibit distinct functions before and after lm: reserve accumulation and energy production, respectively, with both functions dominant in lm-high-abundance mRNAs (Fig. 3). Third, two modules are preferentially active in lm: M7 and M8 may be grouped with modules active during reserve accumulation and desiccation, respectively (Fig. 6). Major seed regulators *LEC1* and *ABI3* are known to interact with diverse genes in mm (Jo et al., 2020). However, in the subsequent lm and desiccation, the TFs and corresponding cis-elements that facilitate the large-scale transition of mRNA populations remain to be determined. Our study may accelerate the discovery of these regulatory components.

The activity of ed-associated-M9 significantly initiates and sustains from early to mid-desiccation, implying a role in the continuum of desiccation processes, which is supported by the functions of TFs within it, as most of them are known to be involved in abiotic stress (Fig. 6; Supplemental Table S10). Remarkably, *GmZAT18* in M9 is featured by becoming highly active in ed and overexpression lines in Arabidopsis show increased tolerance to osmotic stress (Fig.7; Supplemental Fig. S10). ZAT18 is induced by drought and heat stress in somatic tissues in soybean and Arabidopsis (Yin et al., 2017; Wang et al., 2018). Therefore, both endogenous signals and environmental cues may activate *ZAT18*. Whether GmZAT18 and other TFs are regulated by the interaction of ABI5 and chromatin changes, similar to the case during germination, remains to be answered (Wang et al., 2021).

The enriched functions in post-germination are largely different from the stages preceding it (Supplemental Table S3). Meanwhile, some sdlg-enriched functions become dominant in 27hai or even in earlier 16hai (Fig. 4), indicating primary regulatory decisions in sdlg are made during imbibition. These results suggest that most seedling differentiation programs are primarily activated during germination, rather than being encoded in stored mRNAs in dry seeds, consistent with the Harada group’s previous study (Comai and Harada, 1990). However, mRNAs in dry seeds are enriched with hexokinase-dependent signaling, which may initiate regulatory cascades for germination and seedling formation (Supplemental Fig. S6). Furthermore, “nucleosome assembly” functions dominate in germination and include genes encoding histone variants. How these regulatory layers cooperate and which genes in the soybean genome are controlled by these synergistic operations to program germination and seedling growth remain largely unanswered.

The genetic bases and primary regulators of physiological dormancy in Arabidopsis have been extensively studied, such as DOG1 (Iwasaki et al., 2022). By contrast, the molecular bases underlying other dormancy types are relatively unknown, e.g., physical dormancy. The soybean seed coat acts as an impermeable barrier to prevent water absorption and thereby delay germination (Abbo et al., 2014). Recent soybean studies show a seed-coat-expressed gene *GmHs1-1* regulates seed coat composition and water permeability (Sun et al., 2015). While progress has been made in understanding the genetic basis of physical dormancy in the seed coat, the molecular basis of physical dormancy in the embryo cells remains unclear. We found soybean homologs differ from Arabidopsis germination-specific genes in two ways: (i) there are more soybean homologs than Arabidopsis genes, which may confer additive genetic effects, and (ii) soybean homologs are active before germination, from cotyledon development through desiccation and dry stages, unlike the germination-specificity in Arabidopsis genes (Fig. 8; Supplemental Table S11). Additionally, the evolutionary events leading to diverse dormancy types applied by soybean and Arabidopsis are determined by their respective environments. Purifying selection operates on these soybean genes, which highlights the importance of maintaining these genes and their activities in soybean (Fig. 8C). Therefore, we propose a model, as one of the mechanisms, that helps prevent further suppression of cellular activity at the physiological level, contributing to physical dormancy (Fig. 9). The role of this subset of soybean genes in physical dormancy remains to be elucidated.

## Materials and methods

Specific details are contained within Supplemental File 1.

### Plant material

The growth conditions and staging of soybean seeds (Glycine max (L.) cv. Williams 82) are detailed in Supplemental File 1. Seed developmental stages for sampling embryos are determined based on morphological features, including stem, leaves, defoliation, pod, seed length, seed weight, cotyledon, and axis.

### RNA-Seq library construction, sequencing, data processing, analysis, and the identification of coexpression modules

Total RNA was isolated from soybean embryos, using the PureLink Plant RNA Reagent (Thermo Scientific) and treated with RNase-free DNase I (QIAGEN). One ug of total RNA with a minimum RIN (RNA Integrity Number) value of 8 was employed for RNA-Seq library construction using the TruSeq Stranded Total RNA Library Prep Plant kit (Illumina). Hisat2 (Kim et al., 2019) was used to map sequenced reads to the soybean genome (Wm82.a1.v1.1; https://www.soybase.org). The read counts per gene were analyzed using HTSeq (Anders et al., 2015), followed by using DESeq2 (Love et al., 2014) to determine differentially expressed genes. The goseq package (Young et al., 2010) was used for soybean gene GO enrichment analysis with a cutoff FDR (Benjamini–Hochberg multiple testing correction). The WGCNA R package (Zhang and Horvath, 2005) was employed to identify gene coexpression modules of highly correlated expressed genes using DESeq2 normalized data by following published methods (Zhan et al., 2015).

### Sequence divergence analysis

We followed previously published methods (Lin et al., 2010) with modifications (details in Supplemental File 1) to calculate Ka/Ks values for soybean genes compared to Arabidopsis genes.

## Author contributions

Y.-C.W., W.-H.H., C.-J.H., Y.-T.J., J.Z., C.-C.C., T.-F.H., J.-Y.L. designed research; Y.-C.W., W.-H.H., C.-J.H., Y.-T.J., T.-F.H., J.-Y.L. performed research; Y.-C.W., W.-H.H., C.-J.H., J.-Y.L. analyzed data; and J.-Y.L. wrote the paper.

## Acknowledgments

We thank I-Ling Liu for assistance with WGCNA analysis, Yu-Hao Ke for assistance with Arabidopsis experiments, and Dr. Michel Delseny for discussion.

