## Supplemental Figure for "Temporal Transcriptome Analysis Uncovers Regulatory Modules Programming Embryo Development from Embryonic Morphogenesis to Post-Germination"

x: $\log_2$ RNA level; y: $\log_2$ RNA level

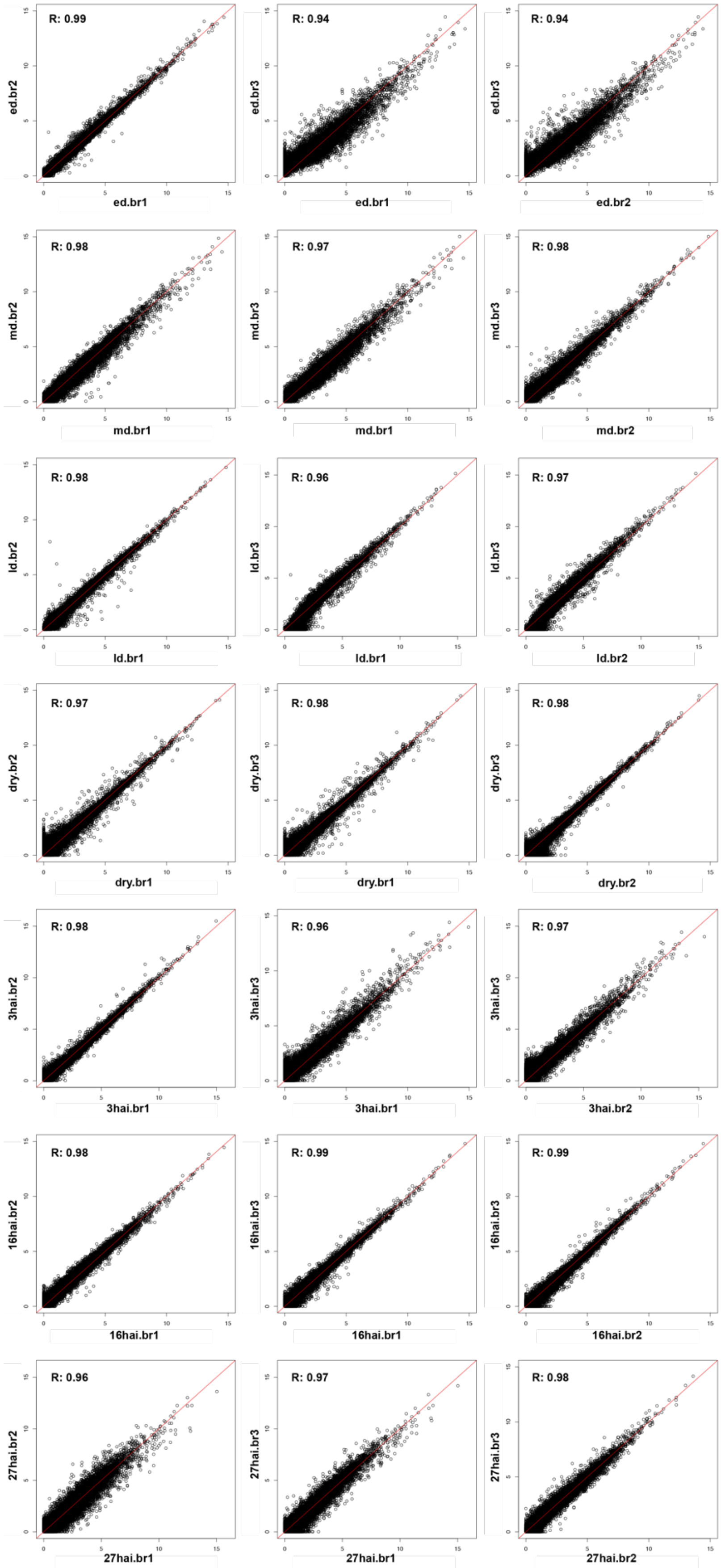

Figure S1. RNA-seq reproducibility of ed, md, ld, dry, 3hai, 16hai and 27hai.
R values were conducted by Pearson correlation coefficient between bio-replicates in the same stage using log2(FPKM +1). Scatter plots were conducted by FPKM of each gene between bio-replicates in the same stage. X- and y-axis is the log2(FPKM +1) of each replicate.

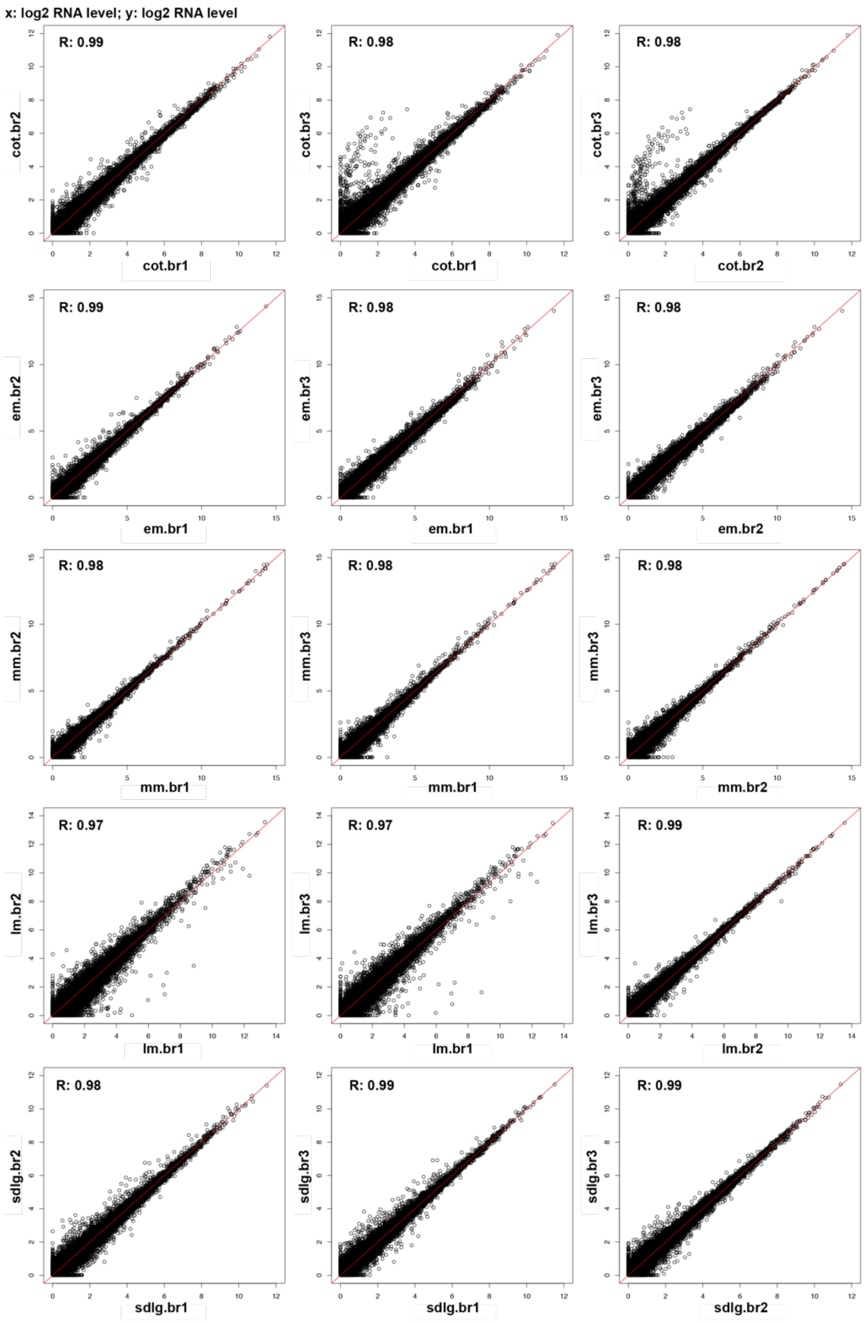


Figure S2. RNA-seq reproducibility of cot, em, mm, lm and sdlg.

R values were conducted by Pearson correlation coefficient between bio-replicates in the same stage using log2(FPKM +1). Scatter plots were conducted by FPKM of each gene between bio-replicates in the same stage. X- and y-axis is the log2(FPKM +1) of each replicate.

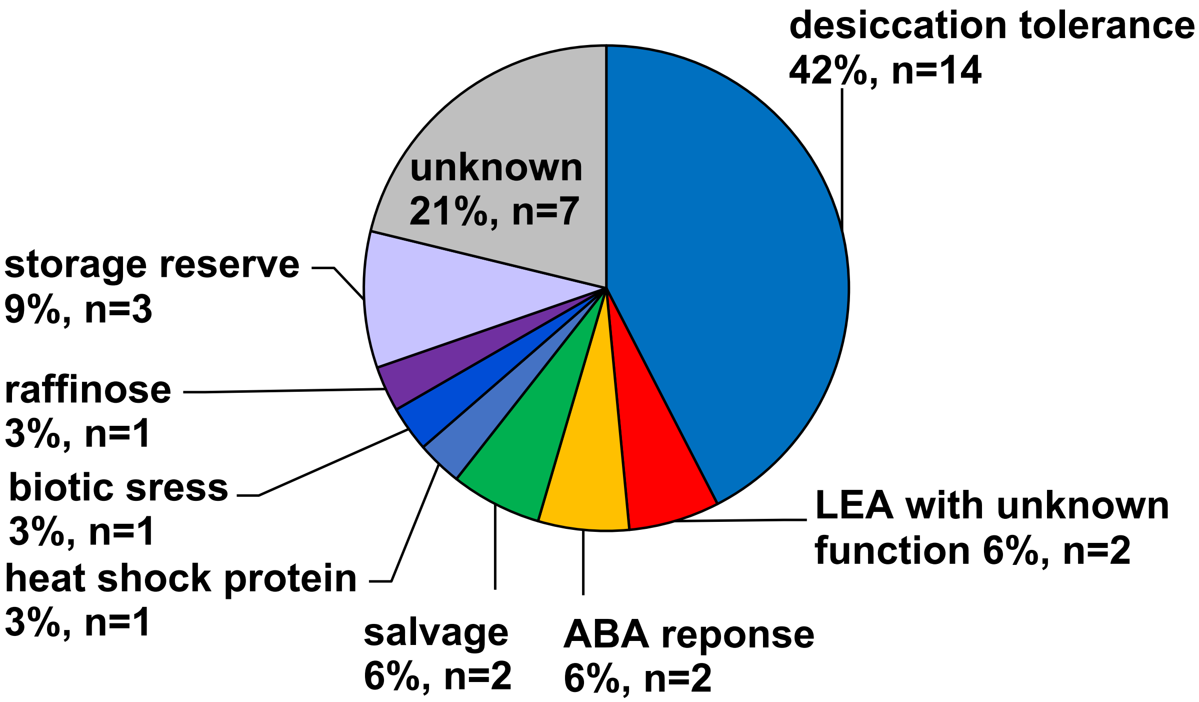


Figure S3. Functional category of high-prevalence genes (> 1,000 FPKM) in the lm.

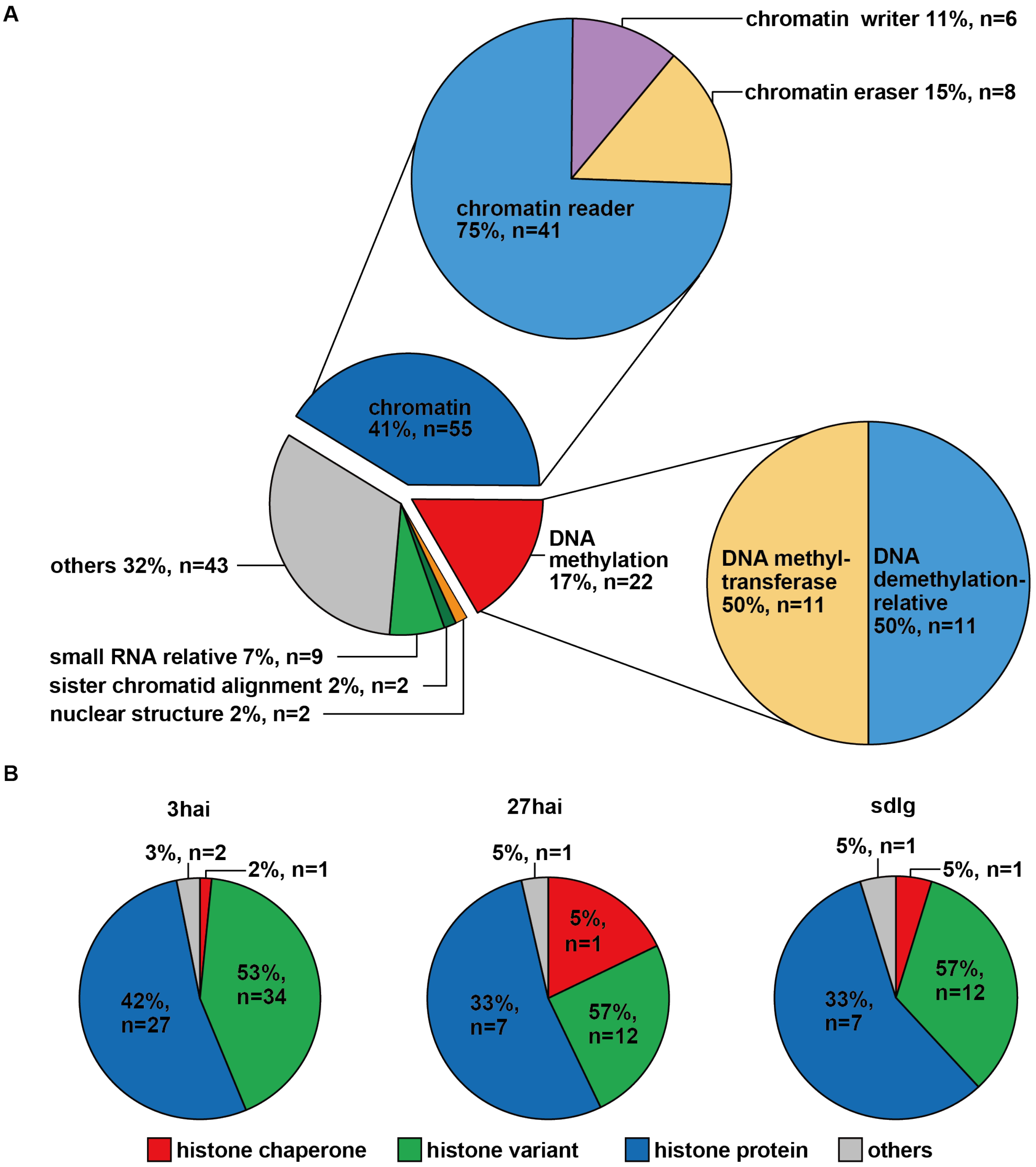


Figure S4. Functional categories of epigenetic-relative GO terms.

(A) Functional category of all significant epigenetic-related GO terms in the mm stage. (B) Functional category of significant epigenetic-related GO terms during germination and post-germination.

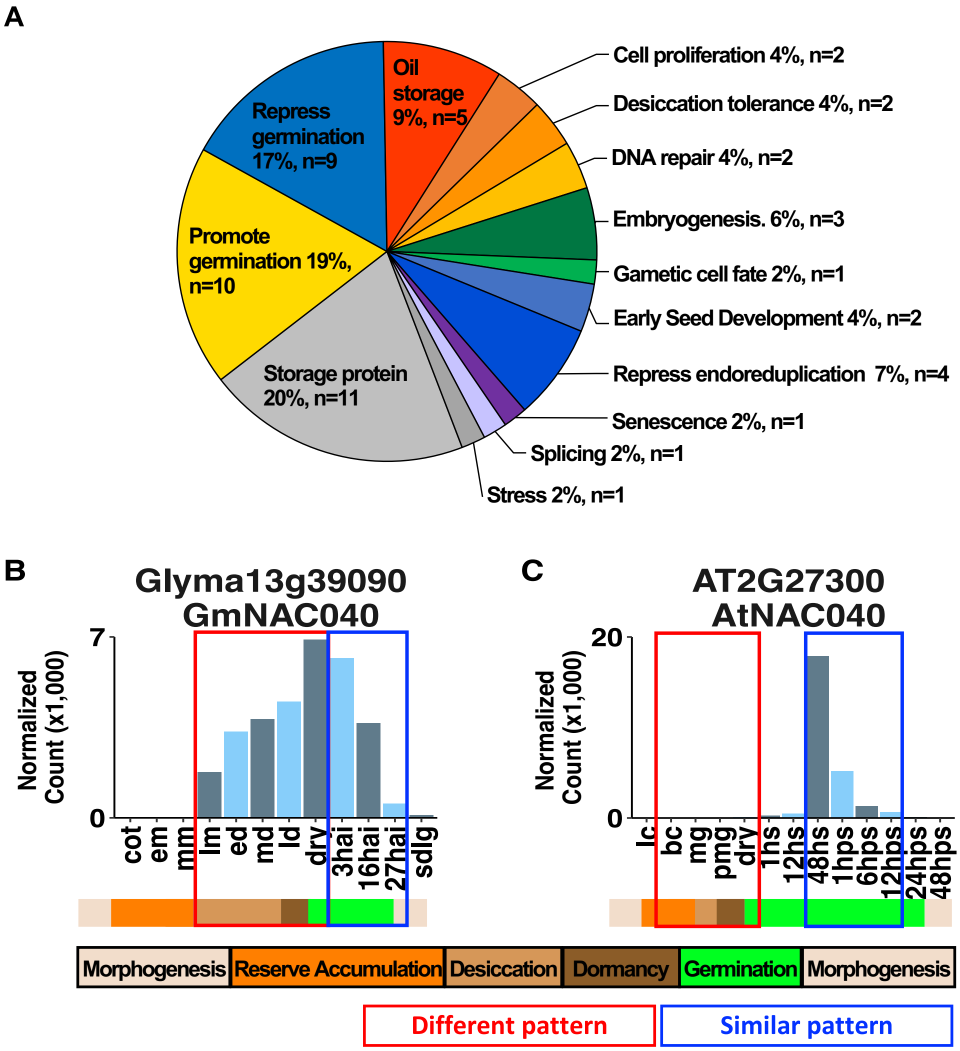


Figure S5. Genes in the GO term "seed germination.

(A) Functional categories of 54 genes in the "seed germination" GO term with gene numbers.

(B) Gene expression patterns of the *NAC040* gene in soybean and *Arabidopsis* are shown in DESeq2 normalized count.

A

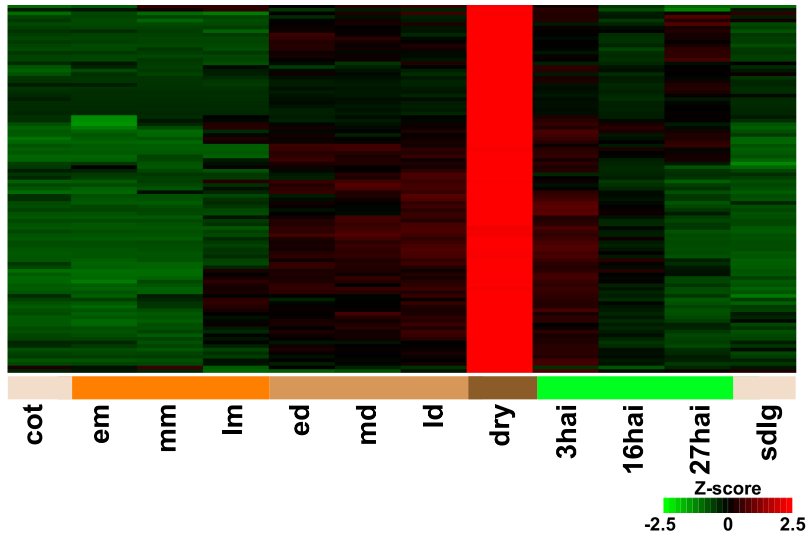


B

| Functional Catalogs | GO ID | GO terms | -log10(FDR) |
|---|---|---|---|
| Cell Death | GO:0012502 | induction of programmed cell death | |
| Energy Related | GO:0006091 | generation of precursor metabolites and energy | |
| | GO:0015979 | photosynthesis | |
| | GO:0019684 | photosynthesis. light reaction | |
| | GO:0009767 | photosynthetic electron transport chain | |
| | GO:0010207 | photosystem II assembly | |
| Respone To Stimulus | GO:0010286 | heat acclimation | |
| | GO:0042542 | response to hydrogen peroxide | |
| Signal Transduction | GO:0009747 | hexokinase-dependent signaling | |
| Transcription | GO:0006354 | transcription elongation. DNA-dependent | |

$-\log_{10}(FDR)$

2 10

Figure S6. Dry-specific up-regulated genes.

(A) Expression patterns of Dry-specific up-regulated genes. (B) Enriched GO terms in dry-specific up-regulated mRNAs.

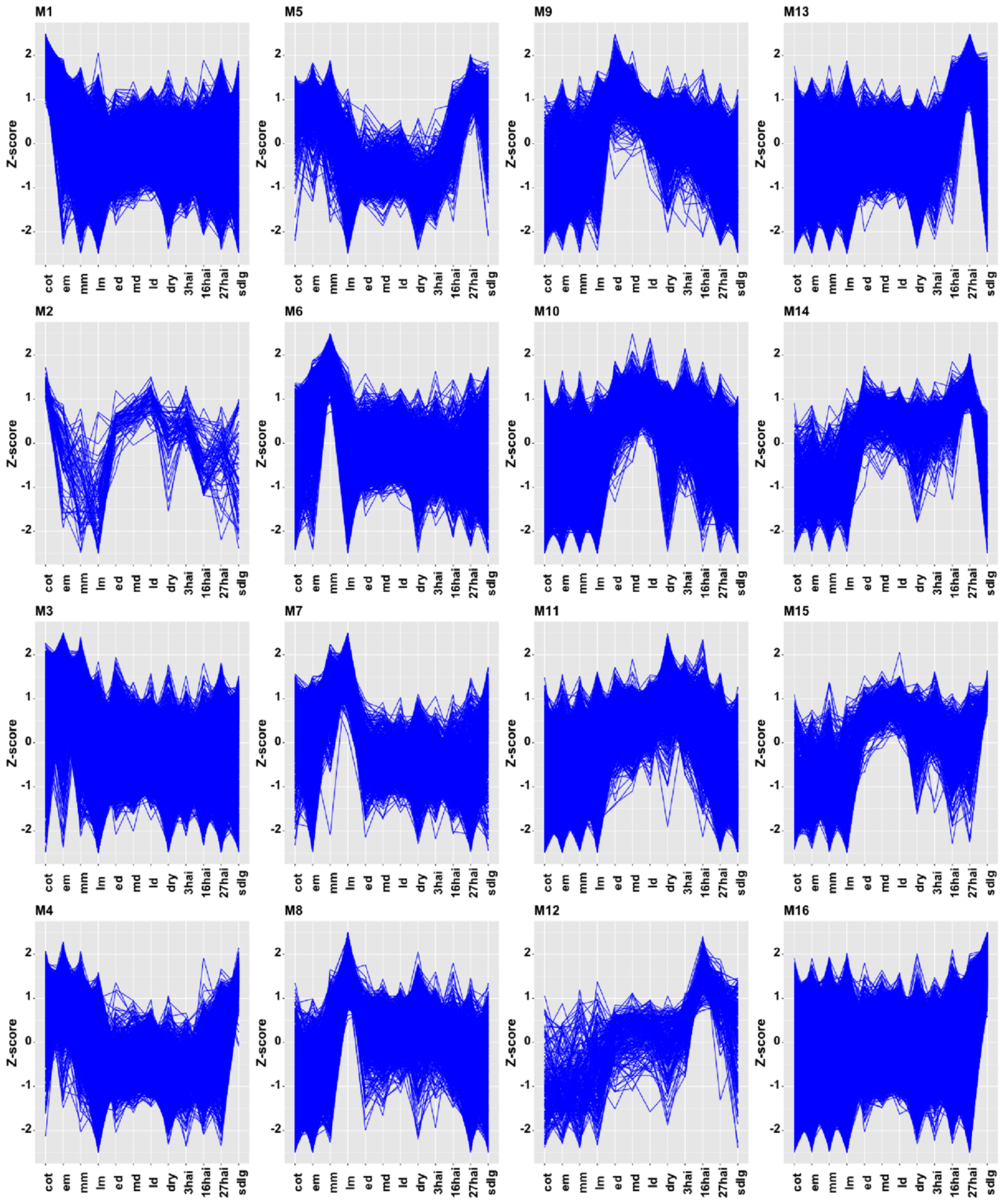


Figure S7. Z-score plots of expression profiles of genes in the module across developmental stages. Trend-plot analysis of the z-scores reveals highly coordinated regulation among genes within the same module.

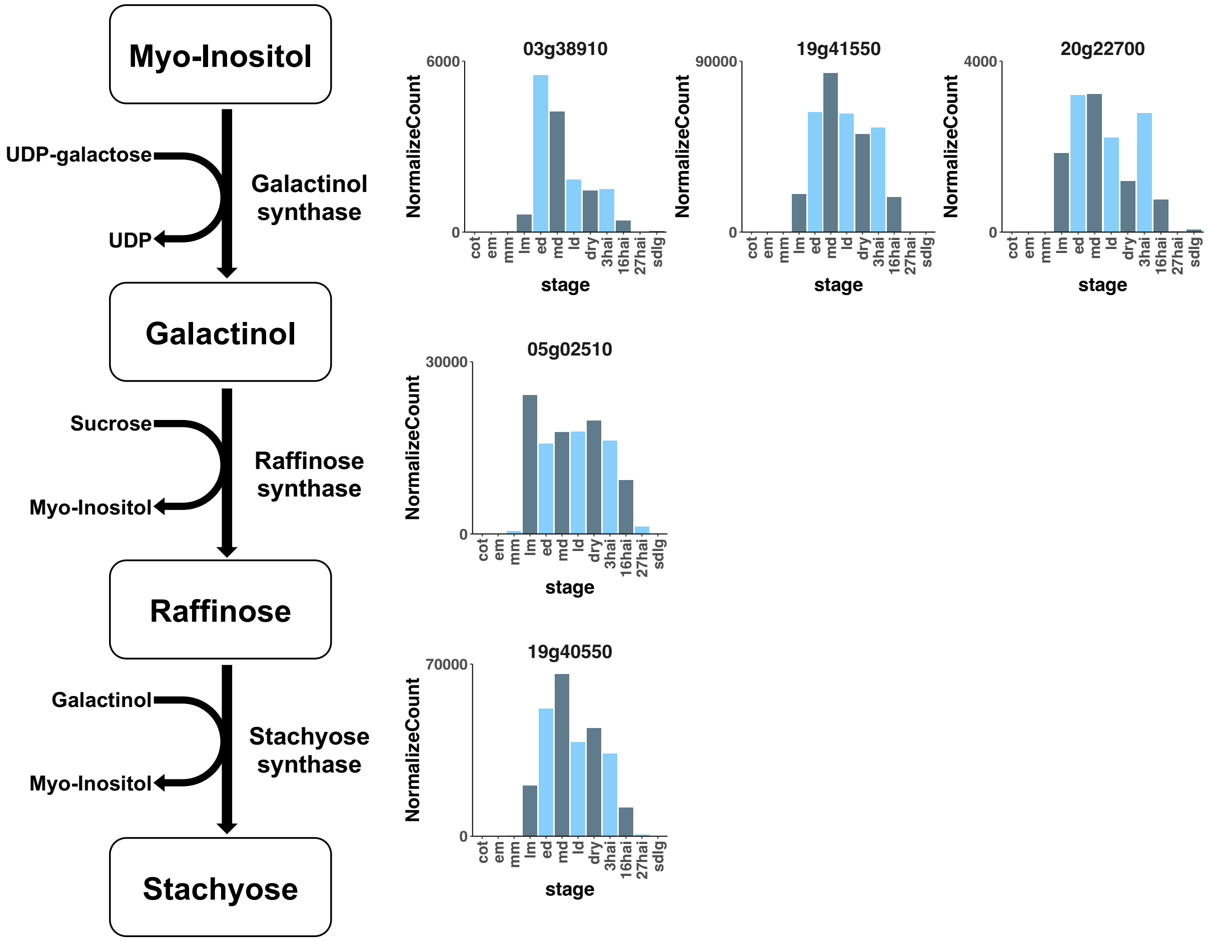


Figure S8. Expression patterns of genes in the biosynthetic pathway of Raffinose family oligosaccharides (RFOs) during seed development.

The RNA level is shown by DESeq2 normalized counts. The pathway is based on published information (Zhuang et al., 2007).

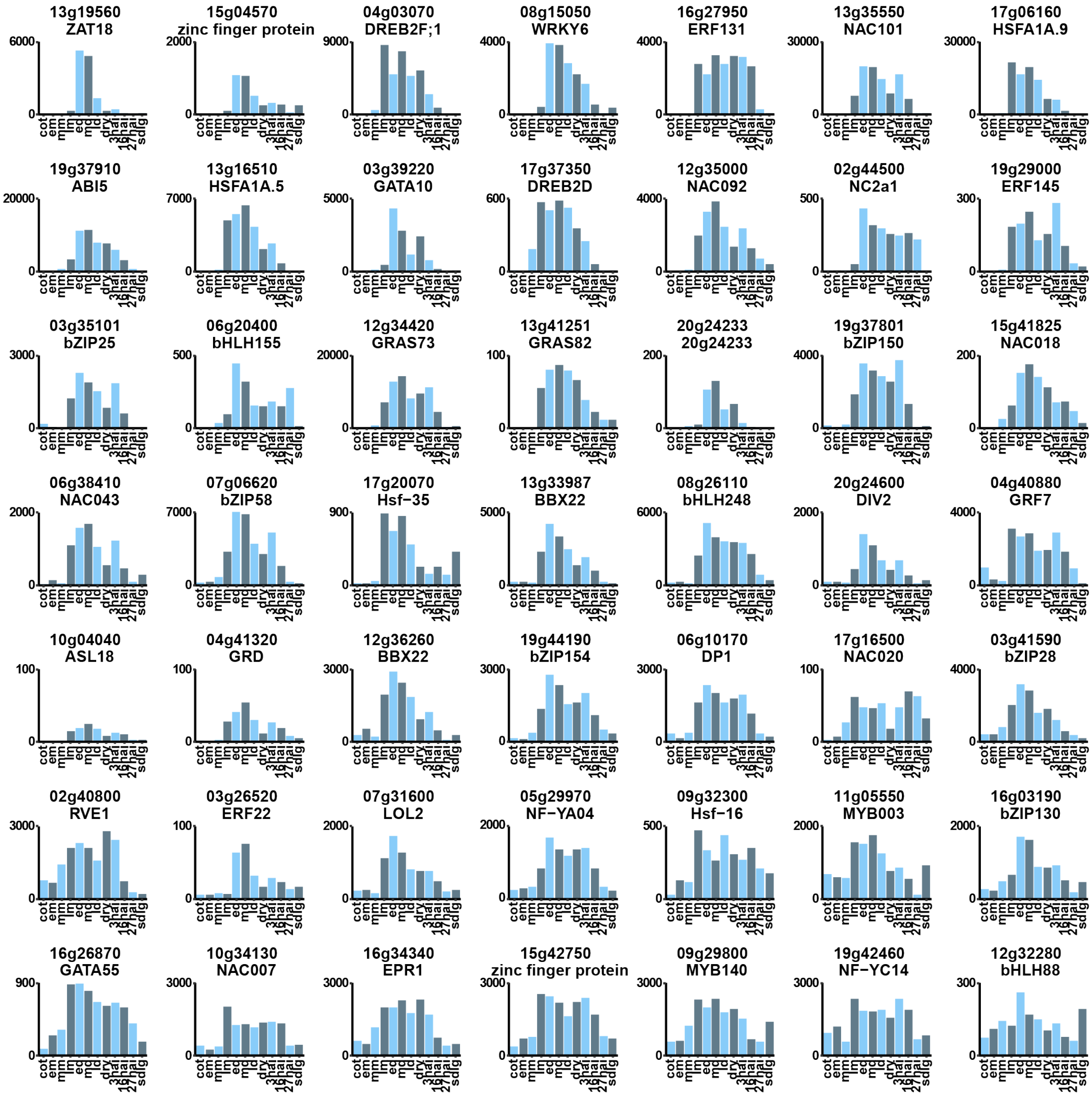


Figure S9. Expression patterns of 49 transcription factor genes in WGCNA module 9.

RNA levels are shown in DESeq2 normalized counts.

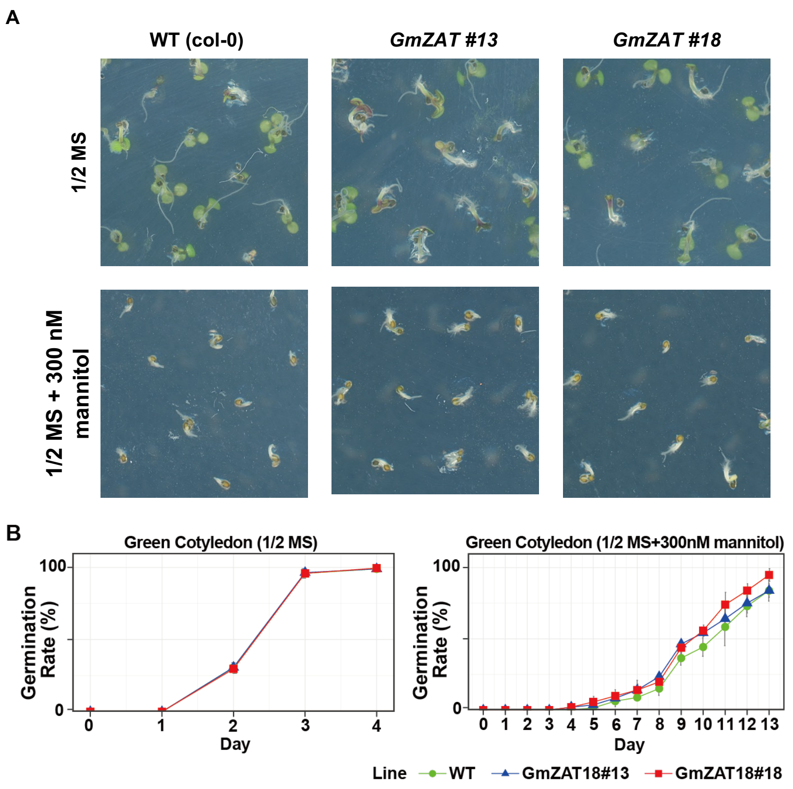


Figure S10. Germination assay of *GmZAT18* overexpression lines under osmotic stress. (A) *GmZAT18* overexpression lines and wild-type plants treated with 300 mM mannitol for 3 days. Germination rates of green cotyledons in GmZAT18 and wild-type plants under control conditions (B) and treatment with 300 mM mannitol (C).

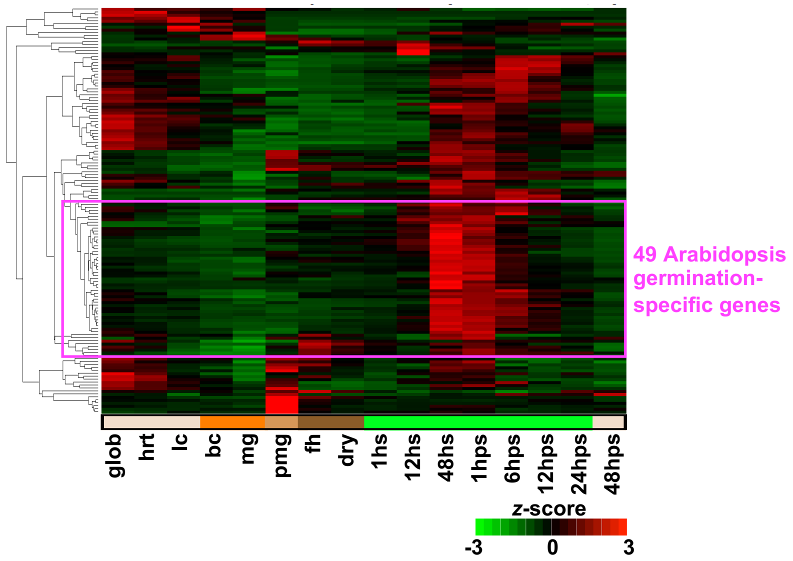


Figure S11. Clustering analysis of 137 *Arabidopsis* genes.

The identification of 49 germination-specific genes in *Arabidopsis* from the RNA-Seq data.
