## Supplemental File for "Temporal Transcriptome Analysis Uncovers Regulatory Modules Programming Embryo Development from Embryonic Morphogenesis to Post-Germination"

### Soybean Growth and seed collection

Soybean plants [*Glycine max* (L.) cv. Williams 82] were grown at 22 °C with under long-day conditions with a 16-h light to 8-h dark cycle in the greenhouse at Academia Sinica-Biotechnology Center in Southern Taiwan. Seeds before germination were staged based on length, weight, and embryonic characteristics, such as shape and color (Supplemental Table below). Drawing of soybean seed embryos at all developmental stages used in our experiments are shown in Fig. 1A. ed, md, ld, and dry seeds had lengths of 10.0-11.0 mm, 9.0-10.0 mm, 8.0-9.0 mm, and 7.0-8.0 mm, respectively. In addition, ed, md, ld, and dry seeds weighed 200-300 mg, 200-300 mg, 150-200 mg, and 100-150 mg, respectively. The leave color, defoliation, stem color and pod color were described in Supplemental Table S1. 3hai, 16hai and 27hai were collected 3 hours, 16 hours, 27 hours after imbibition, respectively. Only embryos were harvested, frozen in liquid nitrogen, ground to a fine powder, and stored at -80 °C before RNA extraction. We collected 3 bio-replicates for each stage.

Supplemental Table. Criteria for staging seeds.

| Stage | Morphology | | | | | | | |
|---|---|---|---|---|---|---|---|---|
| | Stem | Yellow leaves | Defoliation | Pod | Whole Seed Length (mm) | Whole Seed Weight (mg) | Cotyledon | Axis |
| ed | green | 100% | < 50% | green and yellow | 10-11 | 200-300 | yellow, oval shape | yellow |
| md | green | 100% | 50% - 80% | yellow and brown | 9-10 | 200-300 | yellow, round-oval shape | yellow |
| ld | green | 100% | > 80% | yellow and brown, brown, soft | 8-9 | 150-200 | yellow, round shape, not thoroughly hard | yellow |
| dry | green and brown, brown | - | 100% | brown, hard | 7-8 | 100-150 | yellow, round shape, hard | yellow |
| 3hai | - | - | - | - | 8-9 | - | yellow | Wrinkled Seed coat |
| 16hai | - | - | - | - | 10-12 | - | yellow | Axis does not protrude out |
| 27hai | - | - | - | - | 13.5-15 | - | yellow | Axis protrudes up to 3 mm |

### Measurement of seed water content

Newly harvested seeds were dissected to obtain embryos. Thirty-five embryos at the cotyledon stage were used to measure water content. For other stages, ten embryos in each of two replicates were utilized for water content measurement. We recorded the fresh weight of embryos and then subjected them to an 85°C oven for one week for thorough drying. Subsequently, we measured the dry embryo weight for calculating water content using the formula: (fresh weight – dry weight)/(fresh weight). The average fresh embryo weight and water content were then plotted.

**Arabidopsis growth conditions**

The *A. thaliana* Columbia-0 (Col-0) ecotype seeds were sterilized by 50% (v/v) bleach with 0.1% (v/v) Triton X-100, and then washed four times with sterile water. After stratification at 4 °C in the dark for 3 days, seeds were sown on half-strength Murashige and Skoog (MS) medium containing 1% (w/v) sucrose, 0.8% (w/v) agar and incubated in a growth chamber. The growth chamber was controlled at 22°C, 150 μmol photons m−2 s−1, 60% relative humidity and 16-h light/8-h dark cycles.

**Constructs and generation of transgenic plants**

GmZAT18 was synthesized and ligated with pCAMBIA1302 vector to transfer to Agrobacterium tumefaciens GV3101 by floral-dip method. The seeds from transgenic plants were selected on half-strength MS agar medium with hygromycin till we obtained T3 lines.

**RNA extraction and RNA-Seq library construction**

Total RNA was isolated using PureLink Plant RNA Reagent (Thermo Scientific) following the manufacturer's protocol. The extracted total RNA was subsequently purified and treated with DNase using the RNeasy Kit (QIAGEN) and RNase-Free DNase Set (QIAGEN). RNA integrity was assessed using TapeStation (Angilent), and only RNA with a minimum RIN value of 8 (RNA Integrity Number) was utilized for RNA-Seq library construction. One ug of total RNA was employed for RNA-Seq library preparation using the TruSeq Stranded Total RNA Library Prep Plant kit (Illumina).

**Illumina Next-Generation Sequencing**

Paired-end 150-bp reads were generated for the RNA-Seq libraries using Illumina NovaSeq or NextSeq sequencing machines at the sequencing service company.

**Soybean data processing**

Illumina raw reads were processed with Trimmomatic (Bolger et al., 2014) to eliminate adapters and to trim low-quality bases at the 5' and 3' ends, targeting positions with error rates greater than 0.5% and error rate > 0.1%. The trimmed RNA reads were mapped to soybean reference genome Wm82.a1.v1.1 (https://www.soybase.org/) using Hisat2 (Kim et al., 2019). In stages ed, md, ld, dry, 3hai, 16hai and 27hai, pair end mode of Hisat2 was used for mapping forward and reverse strands, and only the uniquely mapped paired reads were used for downstream analysis. In stages cot, em, mm, lm and sdlg 8d, single end mode of Hisat2 was used for mapping and only the reads with mismatch under two and uniquely mapped were used for downstream analysis. Read counts per gene were analyzed by HTSeq (Anders et al., 2015) with soybean annotation.

**Arabidopsis RNA-Seq data in this study**

Arabidopsis Illumina raw reads were downloaded from publicly available datasets:

(i) From global to mature green: glob (globular), hrt (heart; early heart in the download data), lc (early linear cotyledon; early torpedo in the download data), bc (bent cotyledon), mg (mature green) (Hofmann et al., 2019).

(ii) pmg (post-mature green) (Lin et al., 2017).

(iii) From freshly harvested seeds to germination: fh (freshly harvested seed), dry, 1hs (1 hour stratification), 12hs, 48hs, 1hps (1-hour post-stratification), 6hps, 12hps, 24hps, 48hps (Narsai et al., 2017).

**Arabidopsis data processing**

Trimmomatic (Bolger et al., 2014) was applied to assess the base quality of raw reads and trim low-quality bases using the same parameters as those employed for Soybean data processing. Subsequently, the trimmed RNA reads were aligned to the Arabidopsis TAIR10 reference (https://www.Arabidopsis.org) through Hisat2 (Kim et al., 2019). For the stages pmg, fh, dry, 1hs, 12hs, 48hs, 1hps, 6hps, 12hps, 24hps, and 48hps, single-end mode of Hisat2 was employed for mapping, and only uniquely mapped reads were used for downstream analysis. For the stages glob, hrt, lc, bc, and mg, paired-end mode of Hisat2 was used, and only uniquely mapped paired reads were used for downstream analysis. Read counts per gene were analyzed using HTSeq (Anders et al., 2015) with Arabidopsis annotation (Araport11; https://www.Arabidopsis.org).

**Data analysis**

RNA levels of read counts per gene were normalized by FPKM for calculated correlation coefficient between stages and for analyzing high-prevalence genes. Differentially expressed gene analysis was by DESeq2 package (Love et al., 2014) with cutoff q-value $< 0.05$ and $>$ two-fold changes. Hierarchical clustering was analyzed by dChip (Li and Wong, 2003).

**Analysis for germination-specific genes in Arabidopsis**

Narsai and colleagues conducted a microarray analysis (Narsai et al., 2011) to reveal that 775 genes in Arabidopsis are highly active during germination compared to (i) seed developmental stages, including heart, linear cotyledon, bent cotyledon, mature green, (ii) fresh harvest seeds, dry seeds, (iii) seedling, and (iv) somatic tissues, including flower, leaf, and root. Out of the 775 germination-highly-active genes, we focused on 137 genes with mutant phenotypes and/or known subcellular localizations. We fetched publicly available RNA-Seq datasets (see Arabidopsis data processing section above) to examine their gene expression patterns from early seed development (heart stage) through germination and post-germination (48hps). We used dChip for non-supervised hierarchical clustering of these 137 genes to inspect their expression patterns and we found 49 genes exhibit germination specificity (Supplemental Fig. S11). We examined the soybean annotation Wm82.a1.v1.1 (https://www.soybase.org/) to determine 127 soybean genes homologous to the 49 Arabidopsis germination-specific genes (Fig. 8). One of the 127 soybean genes remained inactive throughout the soybean RNA-Seq dataset in this study, while the other 126 active genes were hierarchically clustered using dChip to examine their expression patterns (Fig. 8).

**Identification of soybean genes associated with desiccation**

Gene IDs of desiccation-related genes determined by Righetti and colleagues in *Medicago truncatula* (shown in Supplemental Table S10) (Righetti et al., 2015) were used to retrieve their predicted protein sequences from the Medicago genome database (https://medicago.toulouse.inra.fr/MtrunA17r5.0-ANR). Subsequently, the retrieved Medicago protein sequences served as query sequences to blastp against the predicted soybean protein database (https://phytozome-next.jgi.doe.gov) with a cutoff E-value 1E-10. Soybean desiccation-related genes were identified using the blastp results with the highest E-values.

**Identification of soybean genes associated with longevity**

Soybean genes associated with longevity were determined in three previous studies and we identified soybean genes homologous to longevity-related genes in *M. truncatula* from another study.

(i) A previous study determined soybean longevity-related TFs (shown in Supplemental Table S10) (Pereira Lima et al., 2017) and TF genes with expression profiles that correlate with longevity (P50) (PCC>0.9) were used in this study.

(ii) Genes associated with longevity were identified in QTL hotspot regions in soybean (Zhang et al., 2019).

(iii) Genetic markers associated with longevity were identified in previous soybean QTL study (Dargahi et al., 2014), and their corresponding DNA sequences in the soybean genome were retrieved from SoyBase (https://www.soybase.org). Subsequently, the gene loci overlapping or closest to the DNA sequences of the genetic markers were determined using BEDtools functions, "intersect" or "closest".

(iv) Gene IDs of longevity-related genes determined by previous study in *M. truncatula* (Righetti et al., 2015) were used to retrieve their predicted protein sequences from the Medicago genome database (https://medicago.toulouse.inra.fr/MtrunA17r5.0-ANR). Subsequently, the retrieved Medicago protein sequences served as query sequences to blastp against the predicted soybean protein database Wm82.a1.v1.1 (https://www.soybase.org/) with a cutoff E-value 1E-10. Soybean longevity-related genes were identified using the blastp results with the highest E-values.

**Computational validation of module robustness**

The average Topological Overlap (TO) for each identified coexpression module was computed and compared with the TO average of modules with the same size. These were generated by randomly assigning the 36,383 tested genes to 16 modules. We conducted 100,000 permutations of randomly sampled modules for testing. The observed modules were considered robust when their average TOs showed significantly higher compared to the randomly generated modules.

**Assignment of soybean genes to whole genome duplication events**

We determined the whole genome duplication (WGD) events in which the soybean genes, homologous to Arabidopsis germination-specific genes, originated, using data (the whole genome duplicated protein-enconding genes in the Soybean genome) from a previous study (presented in Supplemental Table S11) (Zhao et al., 2015).

**Sequence Divergence Analysis**

We followed previous published methods with modifications using Arabidopsis homologs (Lin et al., 2010) to calculate Ka/Ks values of soybean genes in this study. Briefly, we used ORF finder (http://www.geneinfinity.org/sms_orffinder.html) to confirm coding sequences of genes. Clustal Omega (https://www.ebi.ac.uk/Tools/msa/clustalo/) was applied to align protein sequences followed by manual inspection. Ka and Ks values were calculated using the codeml program in PAML (Yang, 1997) implemented in PAL2NAL (Suyama et al., 2006).
